# Stochastic Procrastination and Optimal Task Division

**DOI:** 10.64898/2026.09.05.749376

**Authors:** Yonatan Loewenstein, Drazen Prelec, H. Sebastian Seung

## Abstract

Procrastination is widespread, costly, and persistent despite strong intentions to change. We show how procrastination can arise in stationary environments, where there are no deadlines, changing incentives, or new information to justify delay. With hyperbolic discounting, stochastic delay, i.e., acting with a fixed probability each period, emerges as the only self-consistent policy: the only policy that resolves belief–policy inconsistency. We further show that standard reinforcement learning algorithms converge to this Pareto-inefficient solution. We extend the self-consistent model to divisible and continuous-action tasks. In these settings, allowing a voluntary break after completing a subtask never increases, and often reduces, expected delay; enforcing such breaks can reduce delay further, often substantially. Optimal task division equalizes completion probabilities across subtasks, implying that early stages of a larger task should be easier when the goal is to reduce overall time to completion. The model therefore gives a formal rationale and exact quantification for familiar remedies such as task division, starting easy, and enforced breaks, while explaining why procrastination can persist even when intentions, incentives, and information remain unchanged.

## 1 Introduction

### 1.1 The phenomenon and the scope of pure procrastination

Personal testimonials (Akerlof 1991), social science data, and the large market for self-help manuals identify procrastination as a substantial problem in daily life. Repeatedly delaying unpleasant or effortful tasks — canceling an unnecessary subscription, scheduling a medical screening, completing a research paper — is a familiar source of embarrassment and regret. Empirical studies show that most employees and students procrastinate regularly (Steel 2007; Hen et al. 2021), with consequences that include reduced productivity and lower earnings (Nguyen et al. 2013), poorer mental health (Johansson et al. 2023), and the postponement of medical care (Sirois and Pychyl 2016). Notably, over 95% of those who procrastinate express a desire to change (Steel 2007). If so, why do they continue?

Delaying unpleasant tasks can have simple explanations. A person may delay a task because she believes that the immediate cost of acting is too high, because the long-term cost of not acting is not too high, or because future consequences are steeply discounted. In such cases, delay is not irrational, even when the long-run consequences are severe. Delay may simply reflect the agent’s preferences and beliefs at the time of choice. Therefore, it is not obvious that the delay is a problem from a welfare perspective. It becomes a welfare problem if the agent’s beliefs or valuations are biased: if she overestimates the immediate cost of acting, underestimates the future cost of delay, discounts future welfare too steeply (relative to what she would endorse under reflection), or represents future effort as much less costly than present effort (Le Bouc and Pessiglione 2022). When delay arises from biased beliefs or valuations, it may be reduced by correcting mistaken beliefs about the task, clarifying the consequences of delay, or persuading the agent to give greater weight to future welfare (Steel 2012). A different source of delay is stress or anxiety, which may increase the actual subjective cost of acting; in such cases, delay may be reduced by lowering that cost directly (Steel 2012).

Because not all delay is irrational, defining procrastination requires some conceptual care (Chebolu and Dayan 2025). Most researchers emphasize three components: delay of task completion, needlessness, and counterproductiveness (Steel 2007). In economics, “needless and counterproductive” may be translated as “Pareto-inefficient” — harmful to the individual from any temporal vantage point (O’Donoghue and Rabin 1999). Therefore, a central feature of procrastination is regret. Procrastinators often do not merely come to regret their delay later; they may experience the delay as needless and self-defeating even while they are delaying.

Much existing research has examined procrastination in tasks with deadlines, including self-imposed and externally imposed deadlines (Burger et al. 2011; Howell et al. 2006; Bisin and Hyndman 2020), where the looming deadline affects behavior through psychological and normative channels alike. We focus instead on *pure procrastination*: delay in stationary decision environments without a deadline, and with no change in circumstances or information. Such cases isolate the core puzzle of procrastination: delay that persists even when waiting has no instrumental value.

### 1.2 Hyperbolic discounting and the procrastination puzzle

Since the 1950s, economists have known that when the shape of the temporal discounting function is non-exponential, individuals may display temporal inconsistency: an action that appears optimal today when planned for tomorrow may no longer seem desirable once tomorrow arrives (Strotz 1955). Subsequent evidence showed that temporal discounting is often non-exponential, in both animals and humans, and is well described by hyperbolic or quasi-hyperbolic forms (Chung and Herrnstein 1967; Keidel et al. 2021; Kirby and Herrnstein 1995).^1^

A large literature, notably (Ainslie 1975, 1992), has discussed how this hyperbolic discounting-driven temporal inconsistency might explain poor self-control with respect to harmful temptations, smoking, overeating and the like, where small pleasures are front-loaded and larger costs delayed. As we shall show formally in the Results section, procrastination presents the complementary case. Here the costs are front-loaded and the benefits are delayed. Fixing a leaky faucet, scheduling a medical appointment, or writing a paper may be worthwhile overall, yet unattractive at the moment of action. This observation was made in early discussions of hyperbolic discounting (Akerlof 1991; Prelec 1989; Prelec and Loewenstein 1997). The conceptual puzzle is that if the procrastinator prefers postponing her action by just one more day, on the next day she will also prefer postponing it, and also on the day after. Thus, she will never complete the task. But if perpetual delay is the implication, then acting now would be preferable. The difficulty is that in that case, acting tomorrow would be preferable to acting today. “I may, for example, prefer to fix the air-conditioner, TV set or leaky faucet next week instead of now, but faced with a stark once-and-for-all decision, I would choose to fix it now rather than have it, say, permanently broken” (Prelec 1989). The crux of the paradox is policy-belief inconsistency. The agent uses her policy to form beliefs about her future behavior; those beliefs, in turn, determine which policy is optimal today. However, it seems that no policy is “optimal” under the assumption that this optimal policy will be followed.

### 1.3 Prior approaches: naïveté and mixed strategies

In a series of influential papers, O’Donoghue and Rabin (1999, 2001) formulated a canonical dynamic model of procrastination and enriched the hyperbolic account by formalizing misperception, or “naïveté”, about agents’ own time preferences. Naïve procrastinators postpone action by one more day in the sincere belief that they will act tomorrow, only to delay again when tomorrow comes. Naïveté is consistent with behavioral evidence and self-reports (Akerlof 1991); however, by itself it does not explain why procrastination in a stationary setting is often temporary rather than perpetual. Additionally, while people often mispredict their own future motivation, why do they not learn, over time, that these beliefs are inconsistent with reality?

Within the perception-perfect equilibrium framework of O’Donoghue and Rabin (2001), Ericson (2017) shows that a fully sophisticated present-biased agent with no benefit from delay has two classes of equilibrium strategies: cyclical strategies, in which the agent acts every *d*^∗^ + 1 periods, and mixed strategies, in which the agent acts each period with a fixed probability. The mixed-strategy probability in Ericson’s model corresponds to the self-consistent stochastic policy we develop below. Because we restrict attention to stationary Markov strategies on the task state, the cyclical class is excluded from the candidate set, leaving the mixed strategy as the unique self-consistent equilibrium. This restriction is also consistent with Ericson’s own selection of the mixed strategy, on the grounds that cyclical equilibria are not robust to the choice of cycle starting period.^2^

Relative to this prior work, we derive the stochastic policy in a Markov decision setting and show how it can be reached through psychologically plausible deliberation and reinforcement-learning dynamics. We then extend the model to divisible and continuous-action tasks, characterize optimal task partitioning, and show why voluntary breaks are expected to reduce delay, and that externally enforced breaks can outperform voluntary ones. The analysis is related to the multi-selves equilibrium framework we developed for operant matching in Loewenstein et al. (2009).

Conceptually, the analysis proposes a third interpretation of suboptimal behavior, distinct from the cognitive-error and preference-conflict frameworks discussed above. A fully sophisticated agent with accurate beliefs and no opposing preference at the moment of choice can nonetheless be trapped at a Pareto-inefficient equilibrium by the structure of self-consistency itself.^3^ This has implications for the taxonomy of behavioral interventions. Debiasing addresses cognitive error; commitment devices and willpower training address preference conflict; neither addresses an indifferent present self trapped in a self-consistent equilibrium. The class of behaviors that fit this third structure is presumably broader than the no-deadline procrastination problem we study here, but characterizing that broader class is beyond the scope of this paper. Our aim is to make the case for the structure itself, by working out one canonical instance of it in detail.

### 1.4 Procrastination as self-consistent delay

Consider the motivation to fix a leaky faucet. Hyperbolic discounting can generate a situation in which fixing the faucet is better than never fixing it, but fixing it tomorrow is better than fixing it today. The agent’s current motivation therefore depends on what she expects herself to do later. If she expects to fix it soon, she has no reason to act today; if she expects a long delay, she has reason to act immediately. Yet, because there is nothing special about today, the latter expectations are hard to sustain: why should she expect to fix it soon if she has no reason to act now? Why expect a long delay if acting immediately is the better choice? Expectations about future action seem to imply a present action that contradicts those very expectations.

The inconsistency is resolved only if behavior is stochastic: the agent acts with a fixed probability each period until the task is done. If that probability is small, a long delay is expected; if it is large, expected delay is small. At an intermediate value, where the immediate cost of fixing precisely balances the expected cost of further delay, the behavior becomes “self-consistent.” We propose that pure procrastination is the manifestation of such a self-consistent stochastic policy.

This self-consistent policy is not efficient. At the equilibrium probability, the agent is just indifferent between acting now and delaying once more. Thus, the present self gains nothing by delaying, while acting now would benefit all future temporal selves. The inefficiency is therefore not a tradeoff between present and future interests, but a consequence of self-consistent indifference. Once delay has occurred, the agent has reason to regret it, which is a hallmark of procrastination.

### 1.5 Divisible tasks and the design of work

The self-consistent policy described above depends on the immediate cost of acting. This raises a natural question: can procrastination be reduced by changing not the agent’s preferences or beliefs, but the structure of the task itself? This distinction is clearest when the intervention is external. A parent concerned about a child procrastinating on homework can try to change the child’s beliefs or valuations, lower the subjective cost of starting, or alter the structure of the task by dividing the homework into stages and scheduling breaks. Our analysis focuses on this third, structural route. Similar structural interventions may be attempted internally, as self-reframing or self-imposed breaks, but they help only insofar as the agent can implement and sustain them. Otherwise, the proposal collapses back into the original puzzle: why can the agent not simply commit herself to acting tomorrow?

Tasks vary in how easily they can be divided. Some, such as receiving a flu vaccine, have an indivisible final step; others, like folding laundry or washing dishes, are composed of several small subtasks. In particular, some forms of work, like writing this paper (which began more than 10 years ago), can be interrupted almost continuously, after typing each letter. Divisible tasks create natural stopping points, allowing a person to pause after completing part of the work and leave the rest for later. One can also consider *requiring* a break after a certain amount of work is completed. How do these variations influence completion time?

Two competing intuitions arise. Dividing a task could *increase* procrastination: the agent might hesitate to begin, fearing that she will pay part of the cost without obtaining the ultimate benefit of finishing. Alternatively, division could *reduce* procrastination by lowering the immediate cost of action and increasing the likelihood of eventual completion. The question is not only theoretical. Retirement saving, for instance, typically involves a sequence of decisions — selecting a plan provider, choosing an account type, and determining an investment allocation — and a designer who wishes to reduce procrastination must decide whether to bundle these decisions or stage them separately.

We analyze two modes of task division. In the first, breaks are voluntary: the agent may spread the task across days but is also free to continue. In the second, breaks are enforced: after making partial progress, the agent is required to stop and resume later. Our results have three parts.

First, voluntary task division never increases the expected time to completion, and often reduces it appreciably. Second — and more surprisingly — enforcing breaks reduces expected completion time still further, even though mandatory breaks raise the minimum feasible completion time. The intuition is simple: under voluntary breaks, the agent may fear that beginning today will set in motion “all work today,” which deters her from starting at all. Under mandatory breaks, she is guaranteed that today involves only “some work today,” with the rest postponed by construction, making her more willing to begin. The advantage of enforced breaks, however, disappears once the task is easy enough that the agent would complete it deterministically; in that case, enforced breaks merely add dead waiting time. Third, expected completion time is minimized when each subtask, conditional on being reached, is completed with the same probability — a principle of *equal motivation across subtasks*. Under either regime, this implies that early subtasks should be easier than later ones. Together, these results suggest anti-procrastination strategies based on deliberate task partitioning, front-loading easier work, and limiting how much work can be done in a single period.

## 2 Results

We begin with the indivisible task, where stochastic delay emerges as the unique self-consistent policy, and then ask how that policy changes when the same task can be divided. Here we present the formal results under three task-partitioning regimes. Regime 1 is the base case: a single task that must be completed in one period. Regime 2 divides the task into several subtasks with the same total cost, and allows breaks between subtasks. If the agent takes a break, the clock advances by one period. Of particular interest is the limiting, continuous version of Regime 2, obtained by taking the number of subtasks to infinity. The agent can take any number of breaks. Finally, Regime 3 divides the task into finitely many subtasks but, unlike Regime 2, a break is required after each completed subtask.

Our main focus is on expected time to task completion. For each regime and task division, we characterize the induced self-consistent stochastic policy and compare the resulting expected completion time. In this sense, expected time serves as our cardinal measure of intertemporal welfare. We also briefly discuss Pareto welfare comparisons across time periods.

### 2.1 Regime 1: A single, indivisible task

We represent the situation by a Markov Decision Process (MDP) (Sutton et al. 1998), which in the simple one-task case has two states: a leaky-faucet state (*S*) and a terminal fixed-faucet state (*T*, Fig. 1a). Each day the agent decides either to fix the faucet (action *a*) or to do nothing (inaction *n*). Remaining in *S* incurs a daily cost of 1 due to the dripping noise, wasted water, and so on; fixing the faucet incurs this daily cost plus a one-time repair cost *c*. Once fixed, the faucet stays fixed, and the process ends. The agent’s objective is to minimize the expected discounted sum of costs

**Fig. 1:**
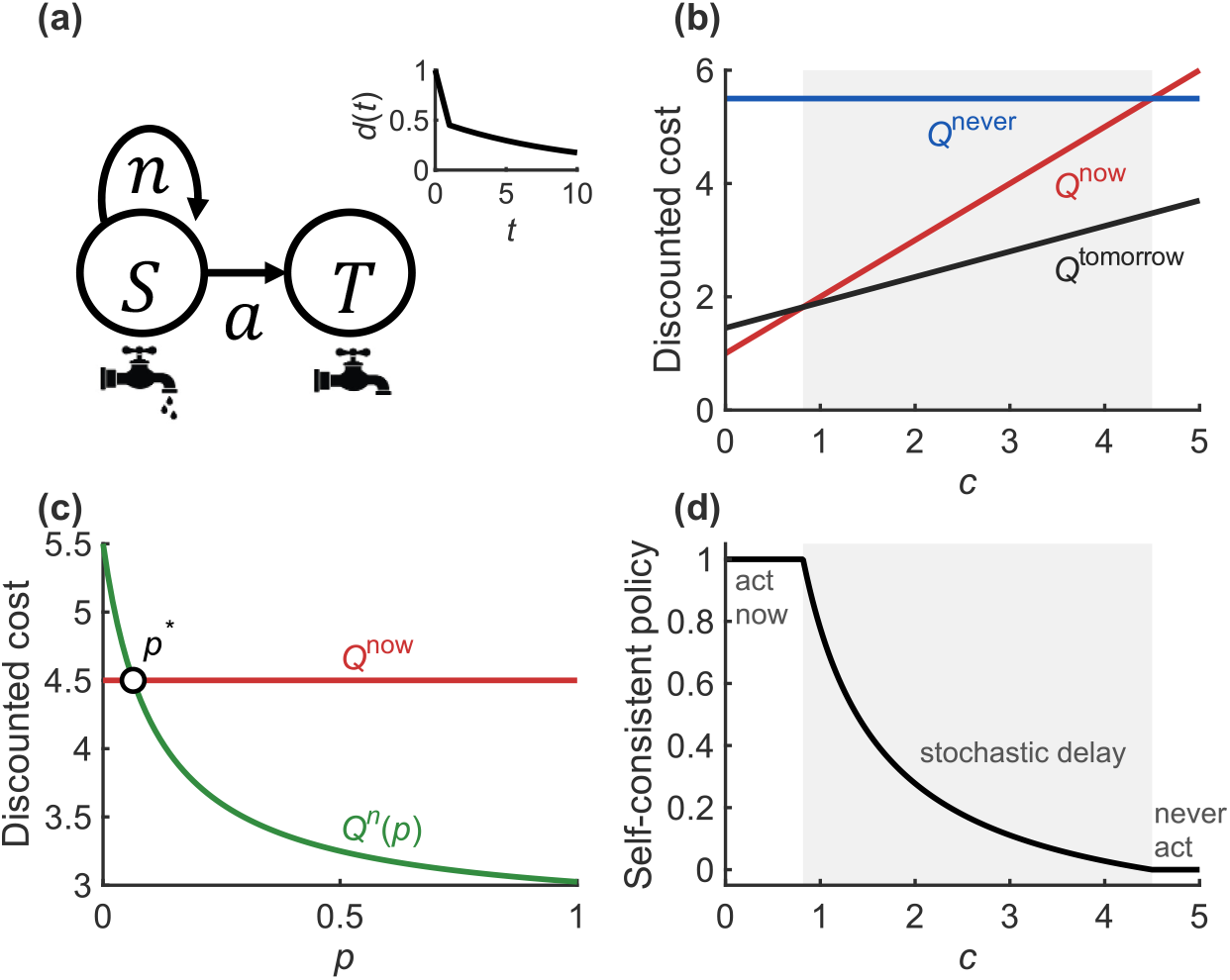
Procrastination in a single indivisible task (Regime 1). (a) Markov decision process for the leaky-faucet problem. The agent starts in state *S* (faucet leaking) and chooses each period between action (*a*) and inaction (*n*). Inaction keeps the system in *S* and incurs a daily cost of 1; action transitions to the terminal state *T* and incurs the daily cost plus a one-time repair cost *c*. The inset shows the quasi-hyperbolic discounting function, with parameters *β* = 0.5 and *δ* = 0.9. (b) Discounted total cost as a function of *c* when the agent fixes immediately (*Q*^now^, red), fixes tomorrow (*Q*^tomorrow^, black), or never fixes (*Q*^never^, blue). For intermediate costs, 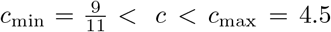 (gray region), delaying by one period dominates immediate action, while infinite delay is dominated. (c) Discounted cost of fixing immediately (*Q*^now^, red) and of inaction (*Q*^*n*^(*p*), green), assuming a fixed action probability *p*. Their intersection (circle) defines the unique self-consistent stochastic policy *p*^∗^. (d) Self-consistent policy *p*^∗^ as a function of *c*. The policy is stochastic in the intermediate-cost region (gray).

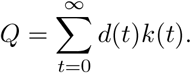

Here *t* denotes the time period measured from the current state *t* = 0, and *k*(*t*) is the cost incurred in period *t*: 1 on a day in which the faucet is leaking and not fixed, 1 + *c* on the day in which it is fixed, and 0 thereafter. The function *d*(*t*) is the temporal discount function. Its decay with *t* determines the relative weight given to future costs. We model time discounting with the quasi-hyperbolic (*β*-*δ*) function (Fig. 1a, inset) (Laibson 1997):

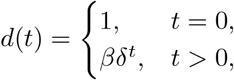

where 0 ≤ *β* ≤ 1 and 0 *< δ <* 1. When *β* = 1, discounting is exponential, with *δ* as the per-period discount factor. When *β <* 1, the agent exhibits present bias, placing extra weight on immediate costs relative to future costs.

Like all discrete-time discounting models, QH partitions continuous time into intervals, e.g., days or weeks. Outcomes falling in the same interval are then assumed to be discounted by the same amount regardless of where they fall within the interval. For example, all events in the current period (*t* = 0) receive weight 1.

According to the QH model, the discounted cost of acting now is

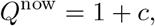

whereas the discounted cost of never acting is

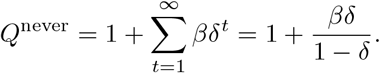

These costs are plotted as functions of *c* in Fig. 1b, in red and blue, respectively. If the cost of repair is not too large, 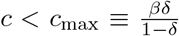, then *Q*^now^ *< Q*^never^, implying that it is better to fix the faucet now than to let it leak indefinitely.

The paradox emerges when considering the discounted cost of acting tomorrow:

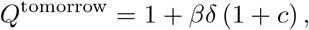

which is plotted in Fig. 1b in black. Comparing *Q*^now^ to *Q*^tomorrow^, if the cost of repair is not too small, 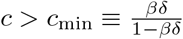, then *Q*^tomorrow^ *< Q*^now^. Thus, it is better to fix the faucet tomorrow than to fix it today.

Thus, for a range of intermediate repair costs, *c*_min_ *< c < c*_max_ (gray in Fig. 1b), the total discounted cost of fixing tomorrow, *Q*^tomorrow^, is lower than the cost of fixing today, *Q*^now^, even though both are smaller than the cost of never fixing, *Q*^never^:

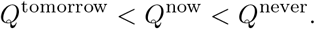

Hence, from the current perspective, it seems best to postpone by just one more day, despite the fact that endless delay is worse. Larger *β* implies a smaller range of intermediate repair costs and under exponential discounting (*β* = 1), the inconsistency disappears because *c*_min_ = *c*_max_ and no values of *c* satisfy both inequalities.

A policy is self-consistent when, conditional on the agent intending to follow it, she has no incentive to deviate. In the intermediate range, no deterministic policy satisfies this condition; we therefore turn to stochastic policies.

Consider an agent whose policy is to fix the faucet with probability *p* in each period. The expected discounted cost of inaction (*n*) under this policy is

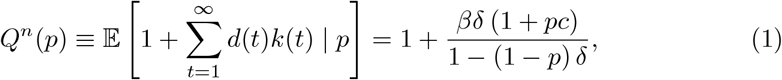

where the expectation is over the stochastic policy *p*.

Figure 1c depicts *Q*^*n*^(*p*) (green) as a function of *p*, together with *Q*^now^ (red). When *Q*^*n*^(*p*) *> Q*^now^, acting immediately is preferable to inaction. When *Q*^*n*^(*p*) *< Q*^now^, inaction is preferred. In both cases, a stochastic policy with action probability *p* is not self-consistent: the agent has a strict incentive either to act or not to act. However, there exists a unique probability *p*^∗^ such that *Q*^*n*^(*p*^∗^) = *Q*^now^ (circle in Fig. 1c):

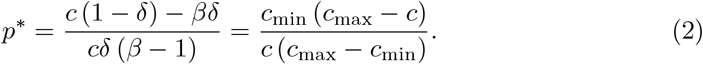

For *c*_min_ *< c < c*_max_, 0 *< p*^∗^ *<* 1. At this value, the agent is indifferent between acting and delaying, and therefore has no incentive to deviate: *p*^∗^ is a self-consistent stochastic policy. When *c < c*_min_, fixing now (*p* = 1) is self-consistent; when *c > c*_max_, never fixing (*p* = 0) is self-consistent. To avoid excess notation, we use *p*^∗^ to denote the self-consistent policy also in these deterministic cases. Figure 1d depicts *p*^∗^ as a function of *c*, showing that the self-consistent policy is stochastic if and only if *c*_min_ *< c < c*_max_.

The same requirement has an equivalent game-theoretic formulation, in which the agent at each time point is represented as a distinct “self.” In this formulation, a self-consistent policy corresponds to a time-invariant (stationary) Nash equilibrium of the game among temporal selves, and such an equilibrium always exists. For *c*_min_ *< c < c*_max_, no deterministic stationary Nash equilibrium exists. The self-consistent stochastic policy *p*^∗^ is therefore the mixed time-invariant Nash equilibrium of the game.

We quantify procrastination by expected time to task completion, which in this model is determined entirely by the task-completion probability. Letting *T* denote the completion-time random variable, for *c*_min_ *< c < c*_max_ we have

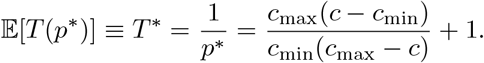

As the repair cost *c* approaches *c*_min_ from within the stochastic region, *T* ^∗^ approaches 1, and the faucet is likely to be fixed immediately. Conversely, as *c* approaches *c*_max_, expected completion time tends to infinity, approaching the never-repair boundary.

### 2.2 Regime 2: Divisible tasks, with breaks allowed

In the Introduction, we raised the question of how task division might affect procrastination, and put forward two competing intuitions. On one hand, dividing the task and allowing a break could increase procrastination: the agent may fear the worst of all worlds, paying the cost of the first part of the task without assurance that the second part will be finished in a timely manner. On the other hand, task division might reduce procrastination if a potentially small amount of effort today would stimulate effort in later periods, making timely completion more likely.

As an example, consider the problem of clearing two pieces of furniture from the basement: a sofa and a table. Rome was not built in a day; Must the basement be cleared in one day, or can it help to divide the work? To analyze this, we extend the model to a two-decision scenario (Fig. 2a). The first decision is whether to clear the sofa (*a*_2_) or do nothing (*n*). If the agent does nothing, she incurs a daily cost of 1 and faces the same decision the next day. If she removes the sofa, she pays a cost *c*_2_ and immediately, in the same day, decides whether to continue and remove the table as well. Choosing to do nothing at this stage again incurs the daily cost of 1 until the table is cleared at an additional cost *c*_1_.^4^ There is no credit for partial work.

**Fig. 2:**
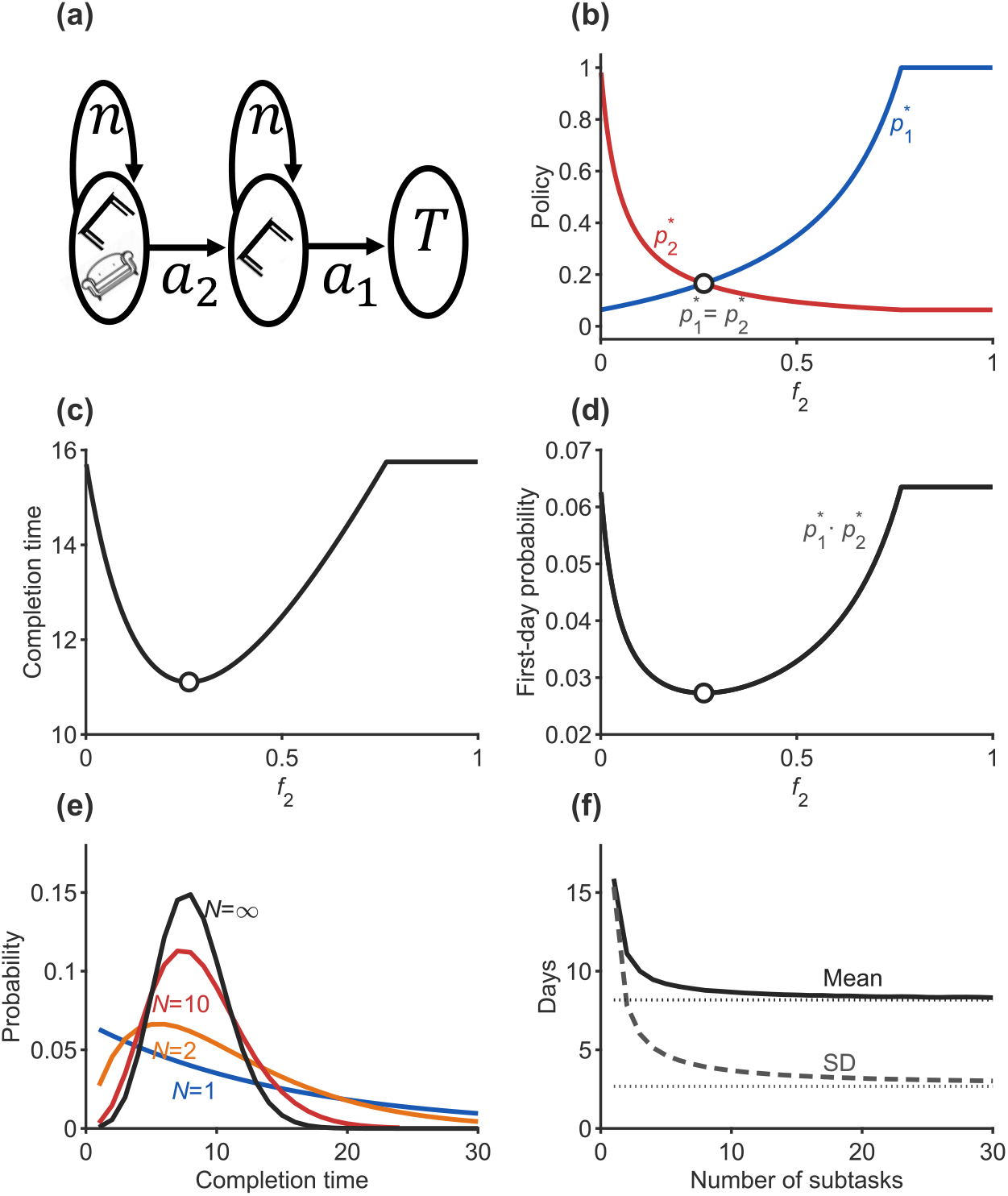
Procrastination in a divisible task with voluntary breaks (Regime 2). (a) Markov decision process for clearing a basement. The agent first chooses whether to remove the sofa (*a*_2_, cost *c*_2_) or delay. After removing the sofa, the agent chooses whether to remove the table (*a*_1_, cost *c*_1_) or delay. A daily cost of 1 is incurred until both subtasks are completed. (b) Self-consistent completion probabilities for the two subtasks as a function of task division, *f*_2_ = *c*_2_*/c* (red: 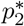; blue: 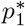). (c) Expected time to task completion as a function of *f*_2_. Completion time is minimized when subtask completion probabilities are equal, 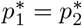. (d) Probability of completing the entire task in the first period, 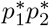. Circles denote the case in which 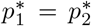. (e) Completion-time distributions for the bundled task (blue), optimal division into two subtasks (orange), optimal division into ten subtasks (red), and the continuous limit (black). (f) Expected completion time (solid line) and standard deviation (dashed line) as functions of the number of subtasks under optimal task division. Dotted lines denote the corresponding continuous-limit values. Parameters: *β* = 0.5, *δ* = 0.9, total cost *c* = 3.5.

We are interested in how task splitting affects procrastination relative to the bundled case, in which both pieces must be removed in the same period if any work is done. We assume that total effort is unchanged by task division: *c* = *c*_1_ + *c*_2_. We thus write *c*_1_ = *f*_1_*c* and *c*_2_ = *f*_2_*c*, where *f*_1_ + *f*_2_ = 1.

The final decision, whether to remove the table after the sofa has already been removed, is equivalent to the leaky-faucet problem with *c* = *c*_1_. We can therefore compute the self-consistent policy for choosing *a*_1_ directly from Regime 1. If *c*_1_ *> c*_max_, the agent never removes the table, even after the sofa has been removed. She therefore has no incentive to remove the sofa, and the task is never completed.

If *c*_1_ *< c*_min_, the agent chooses *a*_1_ with certainty. In that case, when facing the initial decision between *a*_2_ and *n*, under the self-consistent policy, choosing *a*_2_ is followed immediately by *a*_1_. Selecting *a*_2_ therefore leads directly to the terminal state at total cost 1 + *c*_1_ + *c*_2_ = 1 + *c*. Thus, when *c*_1_ is sufficiently small, the divided task behaves like the bundled task, and the self-consistent policy coincides with that of Regime 1.

The nontrivial case arises when *c*_min_ *< c*_1_ *< c*_max_, so that the self-consistent final choice is stochastic. In this case, choosing *a*_2_ leads to a second decision at which *a*_1_ is chosen with probability 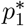, given by Equation (2) with *c*_1_ replacing *c*. The self-consistent policy for the first choice, 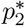, must then make the agent indifferent between choosing *a*_2_ and choosing *n*, given both 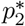 and the downstream policy 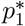.

Within this regime, two cases arise. If the total cost remains too large, *c*_1_ + *c*_2_ *> c*_max_, the task is never completed. Task division therefore cannot induce completion when the agent prefers never completing the task to completing it in a single period. By contrast, if *c*_1_ + *c*_2_ *< c*_max_, the task is eventually completed with probability one. In this case, the self-consistent policy is stochastic (see Materials and Methods).

Figure 2b shows 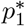 (blue) and 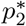 (red) as functions of *f*_2_. When *f*_2_ is small, the initial subtask is relatively easy: 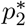 is large, and 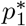 closely resembles 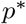, the probability of acting when the two subtasks are bundled together. By contrast, when *f*_2_ is large, 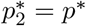 and 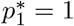. These results allow us to examine how task division, parameterized by *f*_2_, influences procrastination.

The expected time to complete the initial subtask is 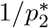, and the expected time to complete the final subtask, conditional on reaching it, is 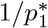. Since both subtasks can be completed in the same period, the expected time to complete the entire task is

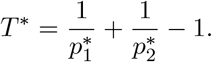

This expectation is plotted in Fig. 2c as a function of *f*_2_. Comparing it with the expected completion time of the bundled task in Regime 1, 1*/p*^∗^, shows that allowing a break never increases expected completion time and often reduces it appreciably. Figure 2c also shows that expected completion time is minimized when the subtask completion probabilities match, 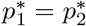 (circle). One can regard this as a principle of “equal motivation across subtasks”: optimal task division makes each subtask, conditional on being reached, equally likely to be completed. This condition implies that the initial subtask must be lighter (*f*_2_ *<* 0.5) because starting carries an extra burden: the agent may pay the first cost without yet obtaining the final benefit.

Notably, although task splitting lowers expected completion time, it also reduces the probability that the entire task is completed on the first day. This probability, 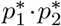, is shown in Fig. 2d as a function of *f*_2_.

These results generalize to tasks divided into more than two subtasks. The self-consistent policy satisfies several properties. First, if the self-consistent policy for the indivisible task in Regime 1 is stochastic, then the self-consistent policy in Regime 2 must involve stochasticity at the subtask level; carving a task into subtasks cannot turn a stochastic indivisible-task policy into a fully deterministic divisible-task policy. Second, stochasticity propagates backward through the task sequence: if the policy is stochastic at a given subtask, then it is stochastic for all preceding subtasks, even when their immediate costs are small. Both properties follow directly from Equations (19) and (20) in the Materials and Methods section.

Third, further dividing a subtask cannot increase expected time to completion (Theorem 1 in the Supplementary Information), but it also cannot increase the probability of completing the entire task on the first day (Theorem 2 in the Supplementary Information).

Finally, when the indivisible-task policy is stochastic, the expected-time-minimizing division equalizes completion probabilities across subtasks (Theorem 3 in the Supplementary Information). The optimal division assigns lower costs to earlier subtasks and higher costs to later subtasks, so that each stage is equally likely to be completed conditional on being reached (Theorem 4 in the Supplementary Information).

It is useful to look at the distributions of time to task completion. For any equal-motivation division into subtasks, the completion-time distribution *T*_*N*_ has a negative binomial distribution shifted by 1. Figure 2e depicts these distributions for the same agent under increasing task divisibility: bundled (blue), 2 subtasks (orange), 10 sub-tasks (red), and the continuous limit (black). First, the mean of the distribution (solid line in Fig. 2f) decreases with the number of subtasks: it is almost 16 days when the task is bundled, drops to almost 11 days when the task is divided into two subtasks, and falls below 8 days when the number of subtasks is large. Second, dividing the task into smaller subtasks eliminates the long tail of the distribution, reducing the standard deviation of completion time (dashed line in Fig. 2f), thereby making completion within a moderate time frame more likely.

As the cost of individual subtasks becomes infinitesimally small, the probability that each subtask is completed under the self-consistent policy converges to 1. However, because the number of subtasks diverges, expected time to task completion converges to a finite value, shown by the dotted line in Fig. 2f. Theorem 5 in the Supplementary Information shows that, as the number of subtasks diverges, the completion time converges to a Poisson random variable shifted by 1, with a rate parameter 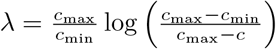.

For the parameters in Fig. 2, the probability of completing the task in the first period is around 6% when the task is bundled, but less than 0.1% as the number of subtasks diverges.

### 2.3 Regime 3: Divisible tasks with mandatory breaks

In the previous section, we showed that task division can mitigate procrastination. The intuition can be seen by recalling the ideal scenario for a hyperbolic agent: “no work today, table and sofa cleared tomorrow.” The task division of the previous section moves partway toward this ideal: the agent can complete a small portion of the work today while leaving the remaining work for a later period, thereby increasing the probability of timely completion. A downside of this policy is that, once the first piece is cleared, nothing prevents the agent from continuing and clearing the second piece as well, producing the less attractive outcome of “all work today.” Starting work today becomes more attractive if the remaining work is guaranteed to be postponed until a later period.

This led us to hypothesize that procrastination might be further reduced by *enforcing* rest periods between subtasks. In this framework, the agent cannot remove the sofa and the table in the same day. If she removes the sofa today, she must take a break and postpone the decision about the table until the next day. Notably, only continuation is restricted: the agent can still do nothing and postpone the decision about the sofa until tomorrow. Trivially, this restriction raises the minimum possible completion time from one day to two. Nevertheless, based on the intuition above, we hypothesized that required rest might, in many cases, reduce expected completion time. From now on, we will refer to the previous regime, in which breaks are allowed but not required, as *libertarian* task division, and to the present regime, in which breaks are mandatory, as *paternalistic* task division.

Again, we look for the self-consistent policy. The second decision, whether to remove the table after the sofa has already been removed, is the same under paternalistic and libertarian task division, since no further break is involved. If *c*_1_ is sufficiently small, the agent chooses *a*_1_ with certainty. For *c*_min_ *< c*_1_ *< c*_max_, she removes the table with probability 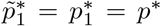. Next consider the initial decision, whether to remove the sofa. Under the paternalistic rule, removing the sofa today leads to the table decision only in the next period, where the table is removed with probability 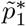. The self-consistent policy for the first choice, 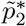, makes the agent indifferent between choosing *a*_2_ and choosing *n*, given both 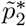 and the downstream policy 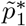.

The results are shown in Figure 3. Figure 3a plots the self-consistent completion probabilities under mandatory breaks as a function of the task division (*f*_2_): 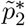 for the initial subtask, shown in red, and 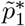 for the final subtask, shown in blue. Because the mandatory-break constraint affects only the transition after completing the initial subtask, the final-subtask policy is unchanged relative to the voluntary-break case, so 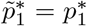. For comparison, Figure 3a also shows the voluntary-break initial-subtask probability 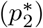 from Figure 2b in light red. As predicted, enforcing a break increases the probability of initiating the task relative to the voluntary-break regime.

**Fig. 3:**
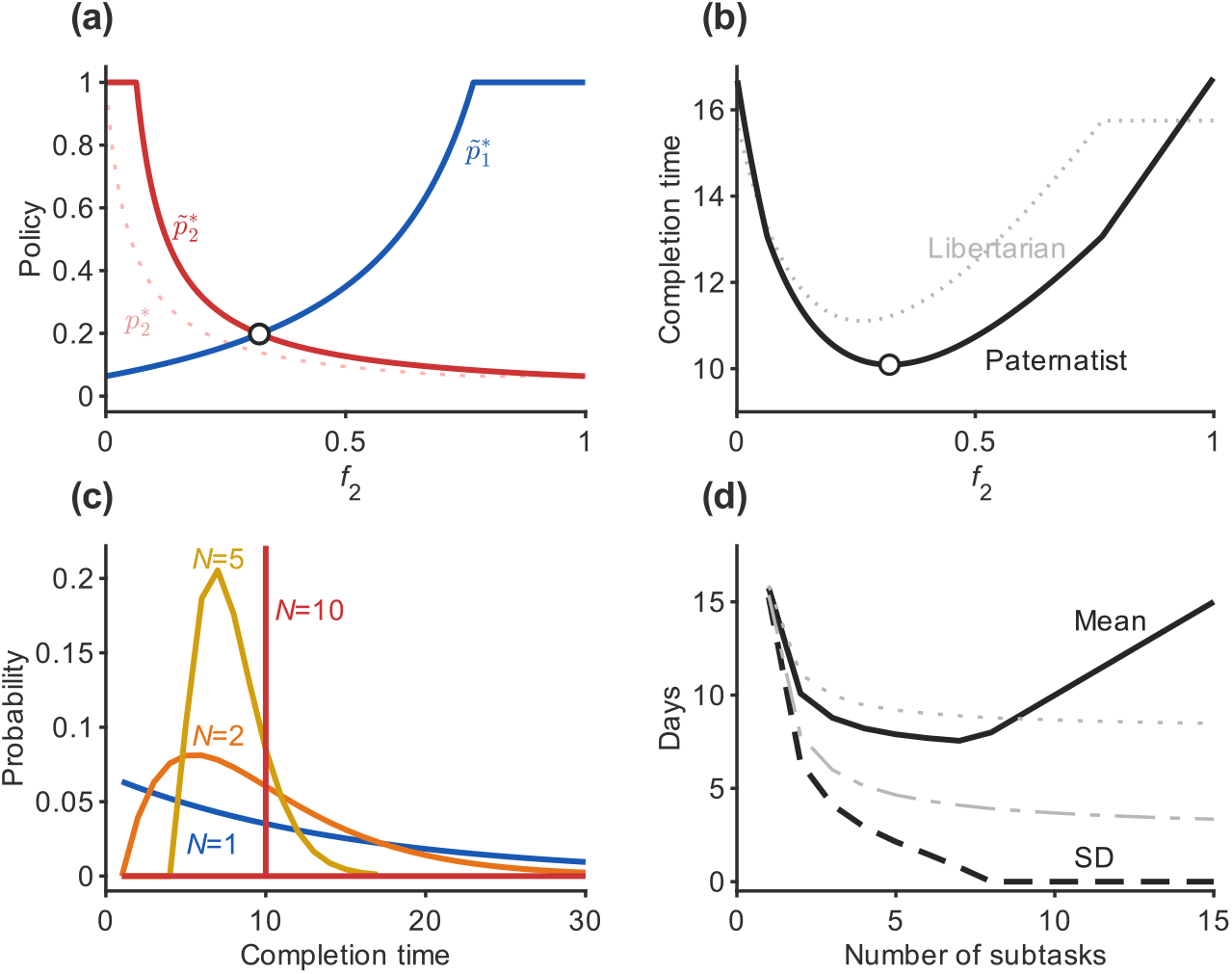
Mandatory breaks and procrastination (Regime 3). (a) Self-consistent completion probabilities under mandatory breaks as a function of task division, *f*_2_ = *c*_2_*/c* (red: 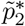; blue: 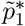). The corresponding initial-subtask probability under voluntary breaks, 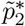, is shown in light red; 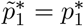. Enforcing a break increases the probability of initiating the task. (b) Expected time to task completion under mandatory breaks (black) compared with voluntary breaks (gray), as a function of *f*_2_. Circles in (a) and (b) denote the task division for which completion probabilities are equal, 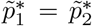. (c) Completion-time distributions under optimal equal-motivation task division with mandatory breaks for the bundled task (*N* = 1, blue), two subtasks (*N* = 2, orange), five subtasks (*N* = 5, yellow), and ten subtasks (*N* = 10, red). For sufficiently large *N*, each subtask is completed deterministically; in that case completion occurs with certainty after exactly *N* periods. (d) Expected completion time (solid line) and standard deviation (dashed line) as functions of the number of subtasks under optimal mandatory-break task division. Thin lines show the corresponding voluntary-break values for comparison. Under mandatory breaks, increasing the number of subtasks initially reduces expected completion time, but once the policy becomes deterministic, further division increases completion time by adding required waiting periods. Parameters: same as in Fig. 2.

Under mandatory breaks, the two subtasks cannot be completed in the same period, so the expected completion time is the sum of the expected waiting times for the two subtasks:

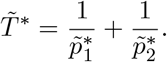

This is shown in black in Fig. 3b. For comparison, the voluntary-break time from Regime 2, *T* ^∗^, is shown in gray. For most values of *f*_2_, the increase in the probability of initiating the task more than compensates for the mandatory waiting period. Thus, although the task can no longer be completed in one day, expected completion time often falls. This comparison is formalized in Theorem 11: whenever the self-consistent policy under mandatory breaks is stochastic, making the break optional, while holding the task division fixed, increases expected time to completion. The benefit of mandatory breaks therefore comes from making initiation more attractive, because starting the task no longer exposes the agent to the possibility of completing all remaining work immediately.

The improvement associated with moving from Regime 2 to Regime 3, by making breaks mandatory rather than optional, generalizes to tasks with more than two subtasks, provided the self-consistent policy under Regime 3 is stochastic (Theorem 7 in the Supplementary Information).

As in the libertarian case, optimal division under mandatory breaks satisfies the principle of equal motivation across subtasks: completion probabilities are equalized across stages. Under this regime, equalization is achieved by assigning larger costs to later subtasks (Theorems 8 and 10 in the Supplementary Information). Consistent with this principle, expected completion time is minimized when *f*_2_ is chosen so that 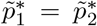 (circles in Figs. 3a and 3b). However, the task division *f*_2_ that equalizes motivation differs between the libertarian and paternalistic regimes.

Mandatory breaks also change the distribution of completion times. Because at most one subtask can be completed in each period, a task divided into *N* subtasks cannot be completed in fewer than *N* periods. For an equal-motivation division, with each subtask completed with probability 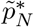, the completion-time distribution is therefore a negative binomial shifted by *N*. Figure 3c depicts these distributions for the bundled task (blue, identical to Fig. 2e), 2 subtasks (orange), 5 subtasks (yellow), and 10 subtasks (red). Unlike in the libertarian case, sufficiently fine paternalistic division eventually makes each subtask deterministic, 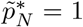. At that point, completion occurs with certainty after exactly *N* periods (e.g., *N* = 10, red).

Figure 3d summarizes the corresponding means and standard deviations. When breaks are mandatory rather than merely allowed, increasing the number of subtasks initially reduces expected completion time, even relative to the libertarian regime. This improvement continues until the subtasks become easy enough to be completed with probability one. Beyond that point, further division is counterproductive, because it only adds mandatory waiting periods. The standard deviation also decreases more rapidly than in the libertarian case and vanishes once completion becomes deterministic.

### 2.4 Learning to procrastinate

An agent need not solve any fixed-point equation in order to behave self-consistently. Starting from any provisional inclination *p* to act today, she samples what would happen if she did not, gauges how costly that would be, and adjusts her inclination accordingly. If inaction appears expensive, she leans toward acting today; if it appears cheap, she leans toward delaying. Iterating this trial-and-error adjustment converges to a unique stable inclination *p*^∗^, at which sampled inaction is exactly as costly, in expectation, as immediate action. That stable point is the self-consistent policy.

To make this precise, let us return to the leaky-faucet problem (Fig. 1a). The agent simulates what would happen if she does not act today, assuming that future choices follow the same inclination *p*. Conditional on inaction today, the simulated realized discounted cost of inaction is

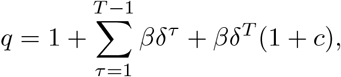

where *T* is the simulated repair time, with *T* = 1 corresponding to repair in the next period. She compares this realized cost with the cost of immediate repair, *Q*^now^ = 1+*c*, and locally adjusts her inclination according to

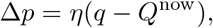

where *η >* 0 is a small adjustment parameter. Note that E[*q*] = *Q*^*n*^(*p*) from Equation (1). Thus, for sufficiently small *η*, stochastic fluctuations average out and the dynamics reduce to deterministic adjustment on the expected costs, with fixed points satisfying *Q*^*n*^(*p*^∗^) = *Q*^now^ — precisely the self-consistency condition that defines *p*^∗^.^5^

This deliberation dynamic also explains why *p*^∗^ is the *unique stable* fixed point in the intermediate cost range. When *p* is near 1, simulated delay is short, so inaction appears cheap and the update pulls *p* downward. When *p* is near 0, simulated delay is long, so inaction appears costly and the update pushes *p* upward. Only the mixed policy *p*^∗^ balances these forces. The same melioration logic underlies our earlier work on operant matching (Loewenstein et al. 2009): a simple local update rule, rather than explicit fixed-point computation, suffices to reach the stochastic equilibrium. The framework extends naturally to divisible-task regimes, where the same updating principle applies locally at each state, with the inclination to initiate or continue work adjusting in response to simulated downstream delay.

The preceding argument describes a deliberative sampling dynamic rather than a reinforcement-learning algorithm: the agent samples possible consequences under a provisional policy and locally adjusts her inclination. We next ask whether the same self-consistent policy can be reached by standard reinforcement-learning algorithms that learn from experienced outcomes.

One way to learn the subjective values *Q*^*n*^(*p*^∗^) and *Q*^now^ is to first learn conventional state values, defined as expected sums of exponentially discounted costs, and then convert these values into quasi-hyperbolic subjective values (see Materials and Methods). Such state values can be learned by incremental one-step prediction-error algorithms, such as temporal-difference (TD) learning.

A computationally simpler way to learn these values is Monte Carlo learning. In each episode, the agent observes the discounted cost trajectory until task completion and uses this realized trajectory as a noisy sample for updating her estimates of the subjective values 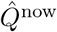 and 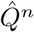.

Finally, the agent can use direct policy-gradient methods such as REINFORCE (Williams 1992) to adjust the policy directly, without explicitly learning the subjective values (Fox and Loewenstein 2025). This is illustrated in Fig. 4. In the simulation, 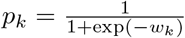, and we use REINFORCE to update *w*_*k*_ from the realized cost *q*_*k*_ and the sampled initial action. The agent in the simulation is initialized such that it begins with a high tendency to act, *p*_1_ = 0.9, and therefore almost always fixes the faucet immediately. Occasionally, however, she postpones the first opportunity and then fixes the faucet soon after. Because such short postponements are associated with lower realized subjective cost than immediate action (bottom), the policy-gradient update gradually reduces the probability of acting (top). Over repeated trials, the agent learns to procrastinate, resulting in longer times to task completion (middle).

**Fig. 4:**
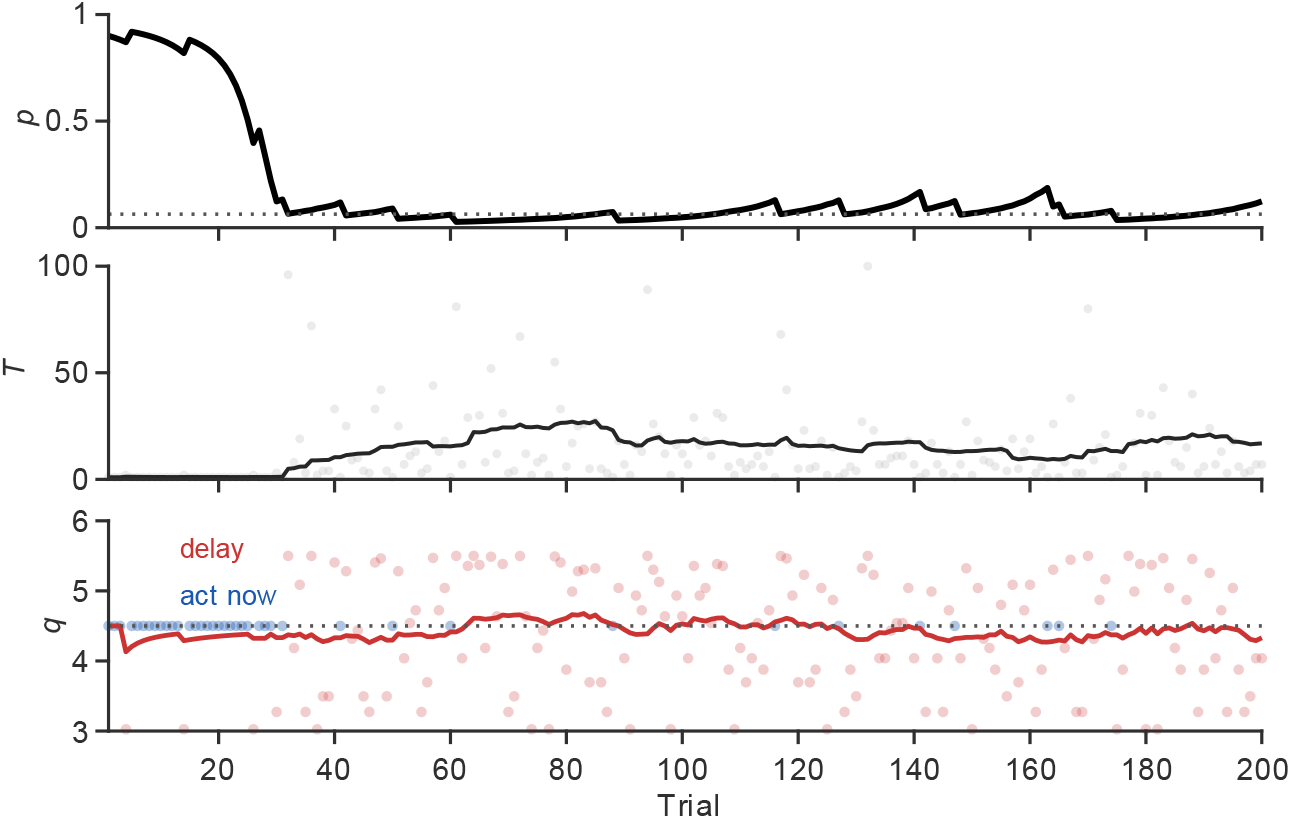
Learning to procrastinate through REINFORCE. The agent begins with a high probability of acting, *p*_0_ = 0.9, and learns from repeated stochastic realizations of the leaky-faucet problem. In each trial *k*, the probability of acting immediately is parameterized as 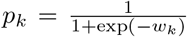. The agent samples an initial action *a*_k_ ∈ { 0, 1}, where *a*_*k*_ = 1 denotes acting immediately and *a*_*k*_ = 0 denotes postponing. After observing the realized discounted cost *q*_*k*_, the policy parameter is updated according to *w*_*k*+1_ = *w*_*k*_ − *ηq*_*k*_(*a*_*k*_ − *p*_*k*_). Top: learned action probability *p*_*k*_, with the dotted line denoting the self-consistent policy *p*^∗^. Middle: realized completion time *T*_*k*_ in each trial. Bottom: realized discounted cost *q*_*k*_, colored by the initial action: blue for acting immediately and red for postponing. Acting immediately yields *q*_*k*_ = *q*^now^ = 1 + *c*, shown by the dotted line; postponing yields a stochastic realized cost determined by the sampled delay. Solid curves are 25-trial causal moving averages.

In all these learning schemes, procrastination emerges as the stable fixed point of a simple trial-and-error process. The agent need not know the parameters of the discount function or compute expected values analytically; through repeated experience she gradually adjusts her inclination until immediate action and expected delay are subjectively balanced. Self-consistent procrastination need not rely on sophisticated foresight or explicit equilibrium reasoning — it can arise naturally as the outcome of standard reinforcement-learning dynamics under quasi-hyperbolic preferences.

### 2.5 Welfare across temporal selves

The analysis so far has described a single agent making repeated decisions over time. As discussed above, the consequences of inaction today depend on the agent’s future actions, which in turn depend on actions further in the future. A standard way to represent these intertemporal dependencies is to treat the sequence of decisions as a dynamic game among temporal selves: each period’s decision is assigned to a different player, or “self.” The time-zero self evaluates costs using today’s discount function; the time-one self evaluates the remaining problem using tomorrow’s discount function; and so on. This framework allows us to evaluate how a policy affects the individual from each temporal vantage point.

We use the standard Pareto criterion across temporal selves (Bernheim and Rangel 2009): strategy *A* Pareto-dominates strategy *B* if every temporal self is at least as well off under *A* as under *B*, with strict improvement for at least one self. Because the model is written in costs, “better off” means lower subjective discounted cost. Under this criterion, the self-consistent stochastic policy *p*^∗^ in the bundled task is Pareto-inefficient. Any self who faces the unfinished task under *p*^∗^ is, by construction, indifferent between acting immediately and delaying once more. But in every history in which the task remains unfinished, that self would have been better off had an earlier self completed the task already. Thus, the deterministic policy “act now” Pareto-dominates the stochastic self-consistent policy, even though “act now” is not itself self-consistent.^6^

Task division complicates this welfare comparison. At first glance, dividing the task might appear to favor the present self, because it reduces the amount of work that must be completed immediately and, under mandatory breaks, may even rule out same-day completion. However, under a self-consistent stochastic policy, the present self is indifferent among all actions that are assigned positive probability. Thus, task division does not improve welfare by making the time-zero self strictly better off. Its main welfare consequences are downstream: it changes the distribution of unfinished work faced by future selves.

Figure 5 illustrates these welfare consequences by plotting subjective cost over time. Panel (a) compares the bundled task (black) with voluntary task division: two subtasks with voluntary breaks (blue) and the continuous voluntary-break limit (red). Panel (b) compares the bundled task with mandatory task division: two (yellow) and eight (orange) subtasks with required breaks. Lower subjective cost corresponds to higher welfare. For the first few future selves, task division can impose higher subjective cost than bundling, either because only partial progress has been made or because mandatory breaks delay completion. From later time points onward, however, divided tasks dominate the bundled task, as future selves become less likely to inherit the entire unfinished task.

**Fig. 5:**
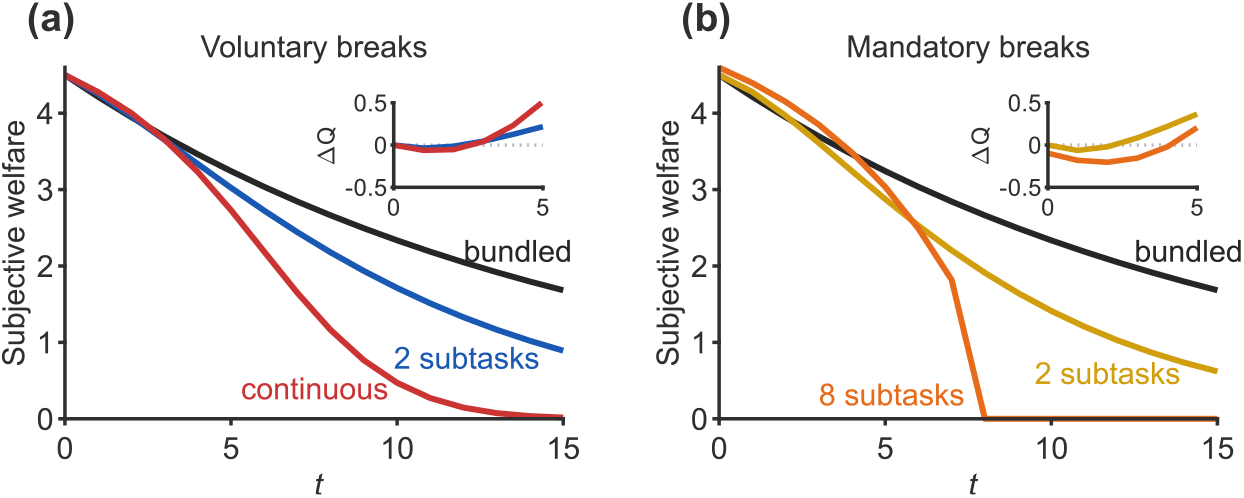
Welfare across temporal selves under voluntary and mandatory task division. Subjective discounted cost is plotted from the perspective of successive temporal selves; lower cost therefore corresponds to higher welfare. (a) Voluntary task division. The bundled task is shown in black, two subtasks in blue, and the continuous voluntary-break limit in red. (b) Mandatory task division. The bundled task is shown in black, two subtasks with mandatory breaks in yellow, and eight subtasks with mandatory breaks in orange. Insets show the difference in subjective cost between the bundled and divided regimes, δ*Q* = *Q*_bundled_ − *Q*_divided_, over the first five periods. Positive values indicate lower subjective cost under task division.

The welfare message is therefore more subtle than the expected-time comparison alone suggests. Task division reduces expected completion time, but it does not uniformly benefit every temporal self at every horizon. Its benefits are primarily realized by later selves, who face a lower probability of inheriting the entire unfinished task. In this sense, anti-procrastination interventions need not make the present self strictly better off in order to be welfare-improving: they may work by changing the distribution of unfinished work faced by future selves.

## 3 Discussion

We have developed a model of pure procrastination: delay in stationary decision environments without deadlines, changing incentives, or new information. Under hyperbolic discounting, delay emerges from a self-consistent stochastic policy, in which the agent acts with a fixed probability each period. We then extended the model to divisible tasks and analyzed how task partitioning influences completion time. Allowing voluntary task division never increases, and often reduces, expected delay, while enforcing breaks further reduces expected time to completion. Optimal task design equalizes completion probabilities across subtasks, implying that early stages should be easier than later ones.

That the only self-consistent policy may be stochastic is a non-trivial observation about behavior. In models of human and animal choice, stochasticity is typically introduced either descriptively, to capture the inherent unpredictability of individual decisions (Shteingart and Loewenstein 2016; Spiliopoulos and Hertwig 2024), or instrumentally, because randomization can support exploration during learning (Fox et al. 2020; Gershman 2018) or strategic mixing in games (Maschler et al. 2020). Yet, exploration is more efficient when directed (and hence deterministic) rather than random (Choshen et al. 2018; Fox et al. 2020), and stochastic choice is common outside strategic settings. Our results identify another source: stochasticity can be required by self-consistency itself. In the procrastination problem, randomization is not added merely as noise, nor justified by exploration or strategic uncertainty; it is the equilibrium form of action when immediate action and expected delay must be made mutually consistent (Loewenstein et al. 2009; Piccione and Rubinstein 1997).

### 3.1 Naïveté as a complementary mechanism

The mechanism developed here is complementary to, rather than competing with, the partial-naïveté account of O’Donoghue and Rabin (1999, 2001). The two accounts identify different mechanisms that can generate the same observable phenomenon: repeated delay despite an intention to complete the task.

In the partial-naïveté account, persistent procrastination is sustained by a systematic mistake about one’s own future willingness to act. The agent each period sincerely intends to act tomorrow, having underestimated the strength of her present bias going forward; tomorrow arrives, the same misestimate recurs, and the task is delayed again. Naïveté is essential to the story: a fully sophisticated agent in this framework, knowing that her future selves will procrastinate too, has nothing to gain from deferring and so acts immediately on a single indivisible task. The procrastination paradox is resolved by introducing the requisite cognitive error.

In the self-consistent stochastic account developed here, persistent procrastination is sustained without any cognitive error. The agent is sophisticated in the relevant sense: she knows that her completion time is geometrically distributed with mean 1*/p*^∗^, and this expectation is, in equilibrium, correct. The paradox is resolved by allowing the policy itself to be stochastic, so that beliefs and behavior can both be self-consistent at *p*^∗^. There is no misestimate to correct.

The two mechanisms therefore yield distinguishable predictions about an agent’s beliefs over her own delay. A naïve procrastinator should be repeatedly surprised at how long the task is taking: her subjective expected completion time should be systematically shorter than the realized completion time, with the gap re-opening each period as she revises her plans. A self-consistent procrastinator should not be surprised in this sense: her subjective expectation of delay should track its realized distribution, even as she regrets it.

The closest existing behavioral test for the naïve–sophisticated distinction is Freeman (2021), who shows that adding an unused extra opportunity to complete a task induces opposite effects on completion time for the two types: a naïf delays further while a sophisticate completes earlier. Freeman’s test cuts between naïveté and sophistication rather than between naïveté and the specific stochastic-equilibrium structure we identify here — the agents in our model are sophisticated in his sense — but the underlying revealed-preference logic, of using completion-time variation across menu manipulations, is the right approach to extend to our finer contrast. The empirical distinction we propose is in principle testable and identifies an empirical edge for the present account. In practice, the two mechanisms likely co-exist, with the relative weight of each varying across individuals and tasks.

The two mechanisms also point to different remedies, whether psychological or institutional. If procrastination is sustained by naïveté, the appropriate intervention is debiasing: better calibration of one’s beliefs about future willingness to act, more accurate forecasts of one’s own behavior, and more reliable feedback about past delays. Naïveté in this sense is one expression of a broader family of biases — over-optimism, planning fallacy, self-deception — that pervade human cognition, and the literature on debiasing addresses procrastination as one symptom of that broader class.

Self-consistent stochastic procrastination, by contrast, is procrastination “with eyes open”: it is sustained even by an agent who forecasts her own behavior accurately, and the problem is not what she believes but what she can implement. The appropriate intervention is therefore not debiasing, but mechanisms of control — devices that allow the present self to bind the future selves. Commitment devices, deadlines, and to some extent, the task division we analyze in Section 2 are all instruments in this second category. The two mechanisms in this sense exhibit the contrast that the literature on intrapersonal welfare has drawn between procrastination as a cognitive failure and procrastination as a self-control failure (Ainslie 1992; Bernheim and Rangel 2009). Notably, the conflict that we study is a conflict between an indifferent present self and unanimous strict preference among future selves.

### 3.2 Remedies

The self-consistent stochastic policy has a simple but important implication for intervention. At the equilibrium probability, the agent is indifferent between acting now and delaying once more. This indifference makes the policy stable, but it also makes it fragile: a small one-time change in the attractiveness of acting today can break the symmetry. The intervention need not permanently change the agent’s preferences, nor should it make the task generally more rewarding. A temporary bonus for acting now, a same-day deadline, or a one-time offer can be sufficient, because it tilts the present choice away from indifference. In fact, it should *not* make the task generally more rewarding. The same bonus for completing the task at some unspecified future time is predicted to be less effective, because it raises the value of acting later as well as acting now, leaving the self-consistency problem largely intact. This logic is familiar in commercial settings. Sellers often use limited-time discounts, expiring offers, or “today only” bonuses to induce immediate purchase. From the seller’s perspective, customer procrastination is costly: a delayed purchase may be forgotten, reconsidered, or lost to a competitor. A temporary incentive for acting now therefore functions as a symmetry-breaking device, shifting the customer from delay to action without necessarily changing the underlying value of the product.

Task design provides a second route for intervention. Two of the central results of the previous sections combine into a concrete prescription for the design of multistage tasks. First, paternalistic enforcement of breaks dominates voluntary breaks in the stochastic regime. Second, optimal task division equalizes completion probabilities across subtasks, which under either regime requires that earlier subtasks be lighter than later ones. Together these results recommend that the designer of a multi-stage process limit how much can be done in any single period, and front-load the easy work, but only to the extent that each stage retains the same self-consistent completion probability.

This prescription stands in interesting tension with two prevailing pieces of advice. The popular “eat the frog first” rule recommends front-loading the *hardest* subtask, on the grounds that completing it removes the largest source of future avoidance. Our analysis suggests the opposite: under hyperbolic discounting and self-consistent stochastic behavior, a heavy initial subtask depresses the probability that the agent will begin at all, and the optimal arrangement is to make the initial subtask easier than what follows. Similarly, the standard advice in choice architecture is to make the first step of any process as small as possible, in order to maximize the probability of initiation. Our results refine this advice: the first step should be small, but only to the point where the equilibrium completion probability of the first stage matches that of subsequent stages.

These design implications are not new as practical advice: they echo familiar recommendations from clinical, educational, and popular approaches to procrastination. Dividing aversive work into smaller steps and beginning with an easy or proximal action is standard in graded task assignment, goal-setting theory, and self-help treatments, where its benefits are often attributed to mastery experiences, increased self-efficacy, reduced task aversion, feedback, or growing confidence (Steel 2012; Fogg 2020). Our model provides a different justification. Even holding beliefs, confidence, mood, and the total task cost fixed, an easy first stage raises the self-consistent probability of initiation because its immediate cost is small while the remaining burden is discounted and assigned to future selves. The optimal easy-to-difficult ordering therefore follows from hyperbolic discounting and self-consistency alone. The model also gives a quantitative criterion for how easy the initial stage should be: it should be calibrated so that its self-consistent completion probability matches that of subsequent stages.

Similarly, scheduled pauses are not new in the popular time-management literature. The Pomodoro Technique, for example, segments work into fixed intervals separated by prescribed breaks (Cirillo 2018). Popular explanations emphasize attention, fatigue, or the subjective manageability of short work bouts. Our framework explains the benefit of binding breaks differently, as a change in the continuation set. When breaks are binding, starting no longer exposes the present self to the possibility of doing all the work today. In the stochastic regime, the resulting increase in the probability of initiation can more than offset the mechanically longer fastest path to completion. Once subtasks are easy enough to be completed deterministically, however, mandatory breaks cease to help and merely add waiting time.

To make the prescription concrete, consider the example raised in the Introduction: enrollment in a retirement saving plan, a setting in which procrastination has been documented empirically as a source of substantial under-saving (O’Donoghue and Rabin 1998). The full enrollment process can be decomposed into selecting a plan provider, choosing the type of investment account, and determining contribution allocations across investment vehicles. We predict greater saving if these tasks are divided into three sequential subtasks. Adding mandatory breaks between them may further increase saving. This contrasts with the common implementation in which the entire enrollment is presented as a single “high friction” decision and the design effort goes into reducing the friction of *every* step uniformly.

### 3.3 Relation to akrasia and choice under indifference

The structural feature just identified — an indifferent present self facing unanimous strict preference among future selves — has antecedents in two literatures that have not previously been brought together. The first is the philosophy of action, where Holton (1999, 2009) departs from the Aristotelian tradition that identifies weakness of will with synchronic conflict between strict preferences (acting against present better judgment) and argues instead that ordinary weakness of will is the failure to maintain a previously formed intention, a failure which need involve no inner conflict at the moment of action. Holton develops this into the positive thesis that intentions function precisely to bridge indifference and incommensurability into action, since desires and beliefs alone do not determine choice in such cases.

The second antecedent is decision-theoretic: the medieval Buridan’s ass thought experiment, in which a rational agent placed equidistant between two equally desirable options has no reason to favor either and starves. Our time-zero self is in just such a Buridan position and resolves it by randomizing with probability *p*^∗^; but where Buridan stipulates the indifference, our model derives it as the unique self-consistent equilibrium of an intertemporal game between successive selves under hyperbolic discounting. The novelty of the present account, properly stated, is the combination: Holton has the philosophical category but no formal model; the Buridan tradition has the randomization but no multi-selves game; the intertemporal-conflict literature has the multi-selves game but presupposes strict-preference conflict throughout. Bringing them together forces the indifferent-present-self equilibrium that the present paper characterizes.

To anticipate one objection, we do not claim that procrastinators can explain their behavior in terms of an explicit stochastic policy. People do not like to think of their actions as random. Subjectively, a decision probability may feel like an inclination or urgency to do the task. Choosing to do the task will depend on that inclination but also on an additional random trigger, which could be external or internal (e.g., (Lebovich et al. 2019)). The agent may attribute the action to a sense that “this is the right day,” just like a player in the zero-sum rock-paper-scissors game may sense that this is “the right moment” to surprise his opponent and play rock. In both cases, de facto stochastic behavior is explained away by pointing to some superficial reason or cue.

The tension produced by the stochastic nature of the self-consistent policy may explain certain negative emotions associated with procrastination. These include regret at not having performed the task earlier and a diminished sense of agency while procrastinating. A procrastinator may appreciate the welfare rationale for acting now and, looking back, regard the delay as a mistake, repeated again and again. Yet she is unable to implement that understanding, as if under a spell. That spell, we conjecture, is the phenomenological experience of action governed by a self-consistent, stochastic policy.

## 4 Materials and Methods

In this section, we provide detailed proofs of the results presented in the main text. For clarity, we repeat some of the equations from the Results section, and the equation numbering continues from that of the main text.

### 4.1 The agent

We consider a QH agent whose objective is to minimize the expected discounted sum of costs:

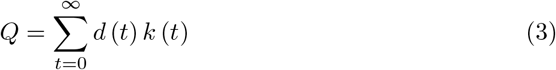

where *t* denotes the time period measured from the current state, *t* = 0, *k*(*t*) is the cost incurred at period *t*, and *d* (*t*) is a QH temporal discounting function:

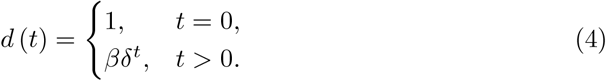

where *β* ≤ 1 and 0 *< δ <* 1 are parameters.

### 4.2 The libertarian task

#### The general framework

We consider a task that can be decomposed into *N* sequential subtasks. For convenience, we index these subtasks in reverse order: the first subtask to be completed is labeled *N*, and the final subtask is labeled 1. Accordingly, in state *i*, the agent must complete subtasks 1 through *i* to finish the entire task. Each subtask *i* carries an associated cost *c*_*i*_ *>* 0, and we assume that the total task cost satisfies

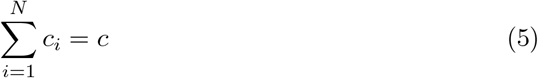

where

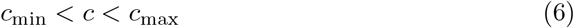

and

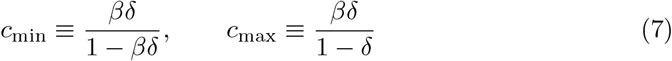

These costs are in addition to a daily cost of +1 before the task is complete.

Note that when 0 *< δ <* 1, a non-empty range satisfying this condition exists if and only if *β <* 1.

In this subsection, we write an expression for the self-consistent stochastic policy as a function of the costs *c*_*i*_ and the parameters of the discounting function *β* and *δ*. This expression will be used in subsequent subsections to study the effect of different task division strategies on average task completion time.

In Reinforcement Learning, the value of a state is typically defined as the expected sum of exponentially discounted rewards obtained when starting from that state. Here, we adopt the same convention but apply it to *costs*: using the discount factor *δ*, we define *V*_*k*_ as the expected sum of exponentially discounted costs associated with state *k*. We refer to *V*_*k*_ as the *value* of state *k*, noting that an agent with an exponential temporal discounting function seeks to *minimize*, rather than maximize, this quantity.

Similar to the Bellman equation, we can express *V*_*k*_ through a recursive relation:

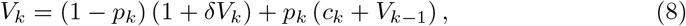

where *p*_*k*_ denotes the probability of acting in state *k*. The first term captures the expected cost of inaction: with probability 1−*p*_*k*_, the agent incurs a daily inaction cost of +1, and remains in state *k*, yielding a future cost of *δV*_*k*_. The second term describes the consequences of completing subtask *k*: the agent pays the immediate cost *c*_*k*_ and transitions to state *k* − 1. Unlike the standard Bellman equation, the term *V*_*k*−1_ is *not* multiplied by *δ*, reflecting that after completing a subtask the agent may immediately proceed to the next one without waiting an additional day. After the final subtask is completed, the agent continues to incur the daily cost of +1 in that day, but not in subsequent days, and therefore *V*_0_ = 1.

Rearranging Equation (8), we can express the action probability *p*_*k*_ directly in terms of the value functions:

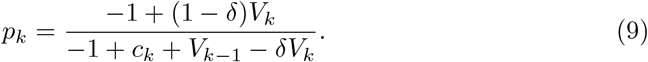

We later evaluate these quantities under the self-consistent policy adopted by the agent.

While an exponentially discounting agent seeks to minimize *V*_*k*_, this is *not* the objective for an agent whose temporal discounting follows the QH formulation. In that case, the value function *V*_*k*_ does not capture the present-bias parametrized by *β*.

We next consider the *subjective* costs of acting and not acting in state *k* for a QH agent, denoted by 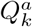 and 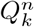, respectively. These quantities represent the expected accumulated costs of the two choices, evaluated under the *β*-*δ* temporal discounting function, and in general, similar to the value function, depend on the probabilities *p*_*k*_. We begin with inaction. The subjective cost of not acting, 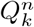, can be written in terms of the value function as

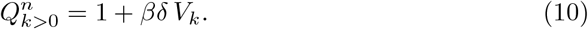

The first term reflects the immediate daily cost of inaction, while the second captures all future costs starting from the next day. Because the “tail” of the *β*-*δ* discount function coincides with the exponential discount function, scaled by *β*, this future contribution is equal to *V*_*k*_ multiplied by *βδ*.

Rearranging Equation (10), we can express *V*_*k*_ in terms of the subjective inaction cost 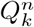:

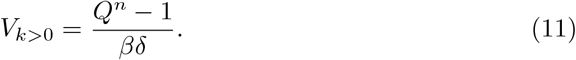

The subjective cost of acting is

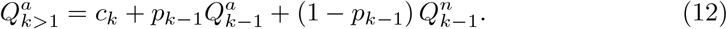

where the first term reflects the immediate cost, while the second and third terms, which depend on the policy *p*_*k*−1_, reflect the subjective cost of state *k* − 1.

For *k* = 1, the subjective cost of acting is simply

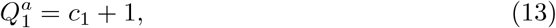

the sum of *c*_1_, the cost of the last subtask and the daily cost +1.

Crucially, under a stochastic self-consistent policy 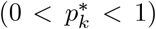, the subjective cost of acting must equal the subjective cost of not acting; otherwise, the agent would deviate from the prescribed policy and choose the option with lower discounted cost. Thus, at a self-consistent stochastic equilibrium,

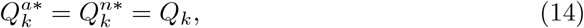

where the asterisk denotes evaluation under the self-consistent policy. Therefore, Equation (12) simply becomes.

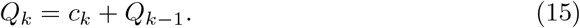

Since 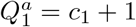, the closed-form expression for *Q*_*k*_ is

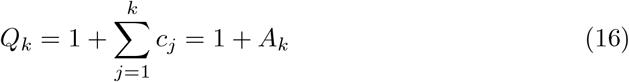

where we define

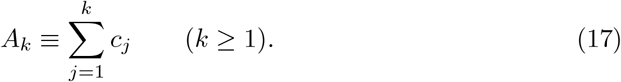

Notably, under a stochastic self-consistent policy 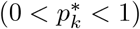, the subjective cost of a state is independent of the discount function parameters *β* and *δ*. Rather, it only depends of the cost of the subtasks left. The mathematical intuition behind this result is that at any state *k*, the subjective values of acting and inacting must be equal, and there is no discounting to the immediate cost of acting.

Substituting Equations (11), (14), and (16) into Equation (9) yields the expression for the self-consistent stochastic policy at each state *k*:

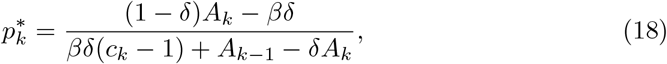

where we define *A*_0_ ≡ *βδ* so that the equation would also hold for *k* = 1 (Equation (2)):

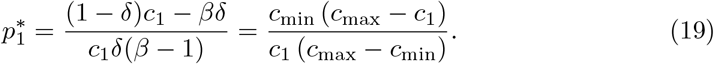

Notably, 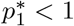 if and only if *c*_min_ *< c*_1_. This is because both 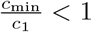 and 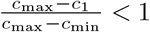 if and only if *c*_min_ *< c*_1_. If *c*_min_ *> c*_1_ then *Q < Q*^tomorrow^ and the only self-consistent policy is to act now. Knowing that the agent will immediately complete subtask 1 in state 1, the consequences of choosing to act in state 2 would be task completion at a cost *c*_2_ + *c*_1_, and so on. Therefore, from now on we will assume that *c*_1_ *> c*_min_.

For *k >* 1, we can write

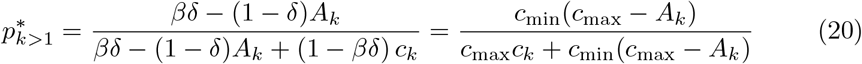

Note that if the task is divided such that the self-consistent solution for the last subtask is stochastic, it will also be stochastic for all other subtasks. To see that, we note that 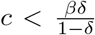. Therefore, *βδ >* (1 − *δ*) *c* ≥ (1 − *δ*) *A*_*k*_. Therefore, according to Equation (20), 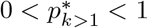.

#### Average time to task completion

A corollary of this analysis is that the average time to complete subtask *k* = 1 is

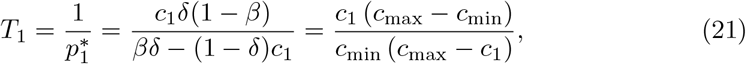

and the average time to complete subtask *k >* 1 is

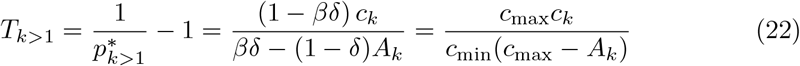

The −1 is added because we consider the action within the larger set of subtasks, and after completing a subtask the agent immediately considers the next subtask.

### 4.3 The paternalist task

#### The general framework

As in the libertarian case, we consider a task that can be decomposed into *N* sequential subtasks. Unlike the libertarian case, at most one action is allowed at each time point. Consequently, the value of state *k* is now given by

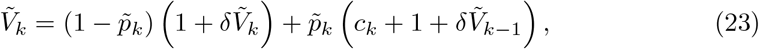

To avoid confusion with the libertarian case, all variables specific to the paternalist task will be denoted by tilde. Specifically, 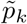 is the probability that the *k*-th subtask will be executed. As before, the first term captures the expected cost of inaction: with probability 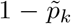, the agent incurs a daily inaction cost of 1, and remains in state *k*, yielding a future cost of 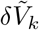. The second term describes the consequences of completing subtask *k*: the agent pays the immediate cost *c*_*k*_ and the daily cost 1 and transitions to state *k* − 1. Unlike Equation (8) and as in the standard Bellman equation, the term 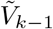 *is* multiplied by *δ*, reflecting that after completing a subtask the agent cannot immediately proceed to the next one. Unlike the libertarian case, 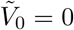. This is because the daily cost of 1 is already included in the cost of action.

Rearranging the equation,

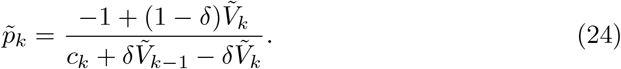

We next consider 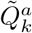 and 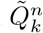, the *subjective* costs of acting and not acting in state *k*, respectively. The subjective cost of not acting is the same as in the libertarian case,

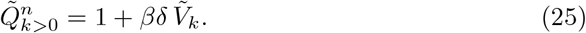

The subjective cost of acting is

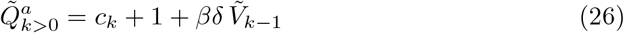

Under a stochastic self-consistent policy 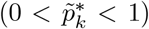, the subjective cost of acting must equal the subjective cost of not acting. Thus, at a self-consistent stochastic equilibrium,

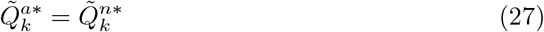

and hence,

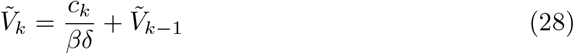

or

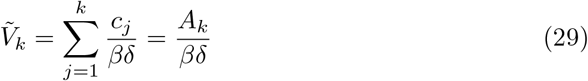

Substituting the result in Equation (26), results in a subjective value that is equal to that of the libertarian case,

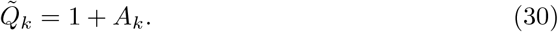

Substituting the result in Equation (24),

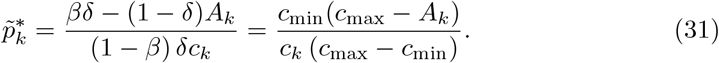

The average time to complete subtask *k* is

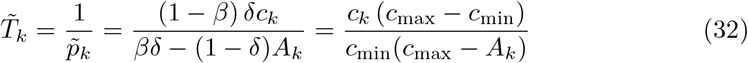

Note that unlike Equation (22), −1 is *not* added to the equation because no more than one subtask is allowed per day.

## 5 Subjective welfare as a function of time

Assume that the task is presented to the agent at time *t* = 0. *In the stochastic regime*, if the remaining task cost at time *t* is *r*, then the subjective value of the corresponding state is 1 + *r*. Thus subjective welfare at time *t* is the expectation of 1+ the remaining task cost.

### Bundled task

When the task is indivisible, the libertarian and paternalist tasks coincide. The agent completes the task each day with probability *p*^∗^, and therefore the completion time is geometrically distributed: Pr(*T > t*) = (1 − *p*^∗^)^*t*^. While the task remains incomplete the remaining cost equals *c*, so

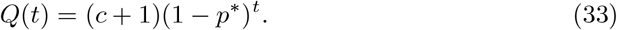

### Two subtasks

Suppose the task is divided into two subtasks with costs *c*_2_ and *c*_1_, where subtask 2 is encountered first and subtask 1 last. Then there are three possible states: state 2: remaining cost *c*_2_ + *c*_1_ = *c*; state 1: remaining cost *c*_1_; state 0: task completed. Let *P*_*i*_(*t*) denote the probability that at time *t* the process is in state *i*. Then

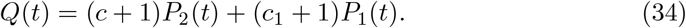

The probabilities of being in each of the states differs between the libertarian and paternalist tasks:

### The libertarian task

Clearly,

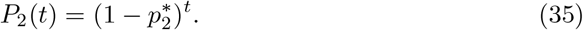

To be in state 1 at time *t*, the first subtask must be completed on some day *s* + 1 ∈ {1, …, *t*}, and the second subtask must then fail to be completed for the remaining *t* − *s* days. Hence

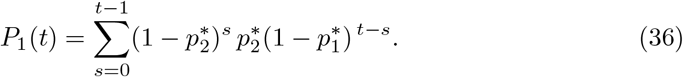

Equivalently, if 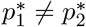,

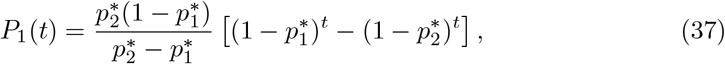

while if 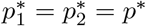,

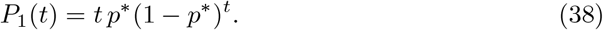

### The paternalist task

The difference from the libertarian case lies in the probabilities of being in each state, because under the paternalist rule at most one subtask can be completed per day.

Let 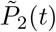 and 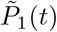 denote the probabilities of the two unfinished states. Then

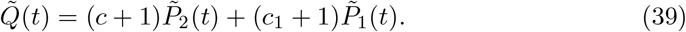

Clearly

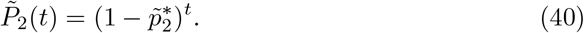

To be in state 1 at time *t*, the first subtask must be completed on some day *s* + 1 ∈ {1, …, *t*} and the second subtask must then fail to be completed during the remaining *t* − 1 − *s* days. Hence

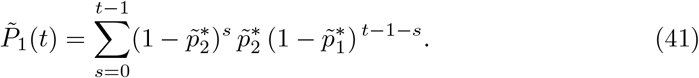

Therefore

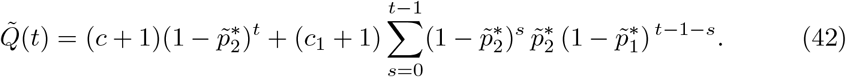

In particular, when the optimal paternalist split equalizes the two completion probabilities, 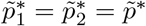, this reduces to

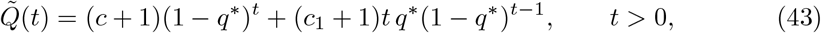

### Continuous task division (The libertarian task)

When the number of subtasks diverges, so does completion time in the paternalist task. Therefore, we will only consider the libertarian task in this limit.

In the continuous limit, the last subtask has cost *c*_min_ and the remaining cost *c* − *c*_min_ is divided into infinitesimal subtasks of constant cost *dc*, with *dc* → 0. Let *x* ∈ [0, *c* − *c*_min_] denote the cumulative completed cost beyond the last subtask. As shown below in Theorem (5), pauses occur along the task according to an inhomogeneous Poisson process in task space with intensity

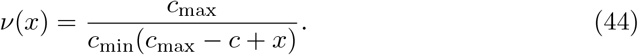

Define the cumulative intensity

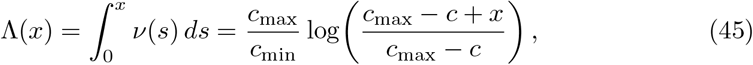

and let

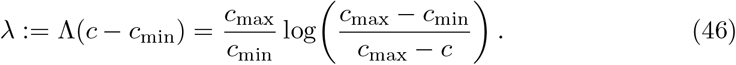

Let *f*_*t*_(*x*) denote the density of the cumulative completed cost after the *t*-th pause. By the standard arrival-time formula for an inhomogeneous Poisson process,

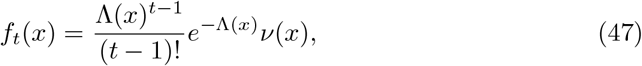

At time *t* = 0, no pause has yet occurred, so *Q*(0) = 1 + *c*.

For *t* ≥ 1, if the task is still unfinished and the cumulative completed cost is *x*, the remaining cost is *c* − *x*. Therefore

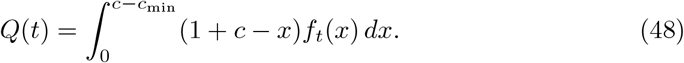

Changing variables to *y* = Λ(*x*), so that *dy* = *ν*(*x*) *dx*, gives

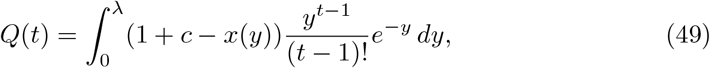

where

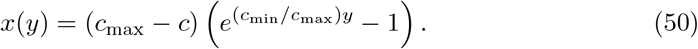

Substituting this expression for *x*(*y*) and evaluating the integrals yields, for *t* ≥ 1,

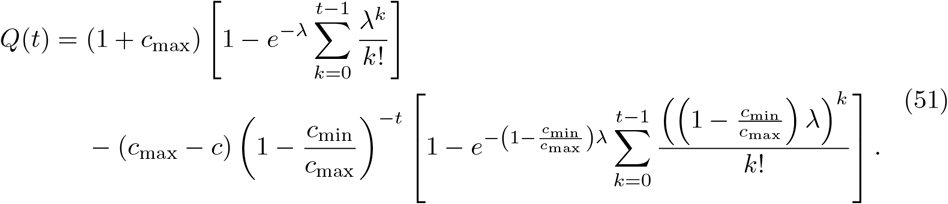

Thus, in the continuous libertarian regime, subjective welfare is obtained by averaging the value of the remaining task over the distribution of progress after *t* pauses.

## 6 Theorems and propositions

### Theorem 1

(Libertarian task splitting weakly reduces expected completion time) *Consider the libertarian regime, and let subtask k have cost c*_*k*_. *Split it into two consecutive subtasks b and a with costs*

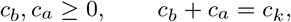

*where b precedes a*.

*Then the expected time contributed by the split subtasks is weakly smaller than that contributed by the original subtask:*

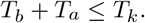

*Consequently, splitting any subtask weakly decreases the expected total time to task completion*.

*The inequality is strict whenever the split subtasks remain in the stochastic regime. In particular, it is strict for every nontrivial split of an interior subtask k >* 1. *For the last subtask k* = 1, *it is strict iff the new final piece satisfies c*_*a*_ *> c*_min_; *if c*_*a*_ ≤ *c*_min_, *then*

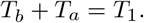

*Proof* We distinguish the cases *k >* 1 and *k* = 1.

**Case 1: splitting an interior subtask** *k >* 1. By Equation (22),

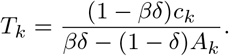

After splitting *c*_*k*_ = *c*_*b*_ + *c*_*a*_, the first new subtask *b* is evaluated at cumulative cost *A*_*k*_, whereas the second new subtask *a* is evaluated at cumulative cost *A*_*k*_ − *c*_*b*_. Hence

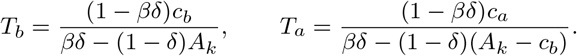

Since *A*_*k*_ − *c*_*b*_ ≤ *A*_*k*_, we have

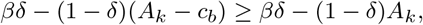

and therefore

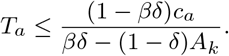

Adding,

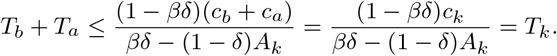

If *c*_*b*_ *>* 0 and *c*_*a*_ *>* 0, then *A*_*k*_ − *c*_*b*_ *< A*_*k*_, so the inequality is strict:

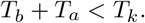

**Case 2: splitting the last subtask** *k* = 1. Before the split,

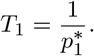

If the new final piece satisfies *c*_*a*_ ≤ *c*_min_, then it is completed with certainty. Therefore, once the agent decides to do *b*, the piece *a* is performed immediately afterward, so the pair (*b, a*) is behaviorally equivalent to the original unsplit last subtask of total cost *c*_1_ = *c*_*b*_ + *c*_*a*_. Hence

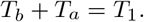

Suppose now that *c*_*a*_ *> c*_min_, so that the new final piece is stochastic. Then by Equation (21),

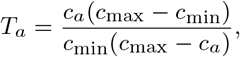

and by Equation (42), since *b* is now an interior subtask with cumulative cost *A*_*b*_ = *c*_*b*_ + *c*_*a*_ = *c*_1_,

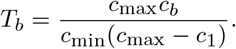

Therefore

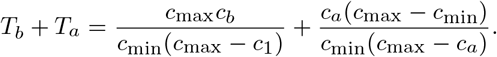

Subtracting from

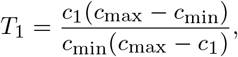

and using *c*_1_ = *c*_*b*_ + *c*_*a*_, we obtain

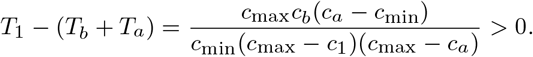

Hence

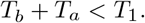

Combining the two cases proves the theorem.

### Theorem 2

(Libertarian task splitting weakly reduces probability of task completion on first day) *Consider an arbitrary libertarian division of a task into sequential subtasks, and let*

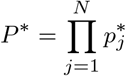

*denote the probability that the entire task is completed on the first day*.

*If one further divides a subtask k into two consecutive subtasks b and a, then the new first-day completion probability* 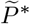 *satisfies*

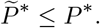

*If k >* 1 *and the split is nontrivial and stochastic, then the inequality is strict:*

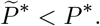

*If k* = 1, *the inequality is strict iff the new final subtask satisfies c*_*a*_ *> c*_min_. *If c*_*a*_ ≤ *c*_min_, *then*

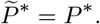

*Proof* Let

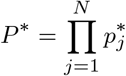

denote the probability that the entire task is completed on the first day under an arbitrary libertarian division into *N* sequential subtasks. Suppose subtask *k* is further divided into two consecutive subtasks *b* and *a*, with

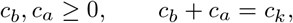

where *b* precedes *a*. Let 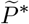 denote the first-day completion probability after the split.

Because, under the libertarian self-consistent policy, the completion probability of a sub-task depends only on its own immediate cost and on the accumulated cost up to that subtask, subdividing subtask *k* does not affect the completion probabilities of the other subtasks. Therefore,

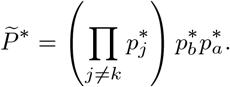

Hence it is enough to compare 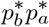 with 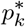. We distinguish two cases.

**Case 1:** *k >* 1. For an interior subtask, we use Equation (20) to write

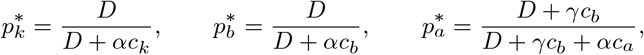

where

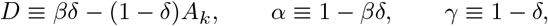

and *c*_*k*_ = *c*_*b*_ + *c*_*a*_. It then computes

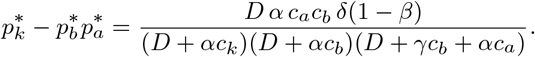

All factors in the numerator and denominator are nonnegative, and for a nontrivial stochastic split they are strictly positive. Therefore

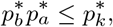

with strict inequality when *c*_*a*_ *>* 0 and *c*_*b*_ *>* 0 in the stochastic regime.

**Case 2:** *k* = 1. Now the split is applied to the last subtask. If the new final piece satisfies *c*_*a*_ ≤ *c*_min_, then the agent completes that final piece with certainty. In that case the split task behaves exactly as if the two pieces were bundled together, so

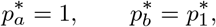

and hence

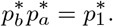

Suppose instead that *c*_*a*_ *> c*_min_, so that the new final piece is stochastic. Then in the bundled case, the probability of completing the task on the first day is (Equation (19))

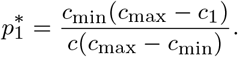

After splitting, the probability of completing the entire task on the first day is (Equation (20))

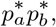

where

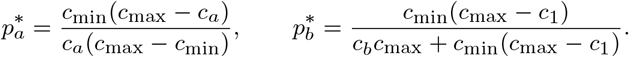

We compare the split and unsplit first-day completion probabilities by taking the ratio:

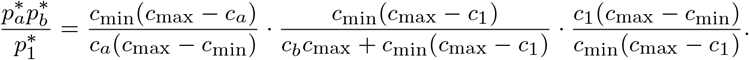

Cancelling common factors yields

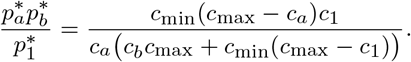

Hence

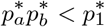

is equivalent to

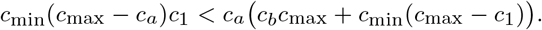

Using *c*_1_ = *c*_*a*_ + *c*_*b*_, the difference between the right-hand side and the left-hand side is *c*_*b*_*c*_max_(*c*_*a*_ − *c*_min_), and by assumption *c*_*a*_ *> c*_min_, this quantity is strictly positive. Therefore

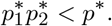

Thus, whenever the newly created final subtask is stochastic, splitting the last subtask strictly lowers the probability that the entire task is completed on the first day.

Combining the two cases yields

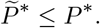

The inequality is strict for every nontrivial stochastic split of an interior subtask, and for a split of the last subtask it is strict exactly when the newly created final piece satisfies *c*_*a*_ *> c*_min_. This proves the theorem.

### Theorem 3

(Optimal libertarian division equalizes completion probabilities) *Assume the libertarian regime and suppose the self-consistent policy is stochastic. Let a task of total cost c be divided into n sequential subtasks with costs c*_*n*_, …, *c*_1_, *where subtask n is encountered first and subtask* 1 *last. Let* 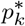 *denote the self-consistent probability of completing subtask k, and let*

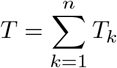

*be the expected time to complete the entire task*.

*If the division minimizes T among all divisions with the same total cost c, then there exists a number* 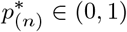 *such that*

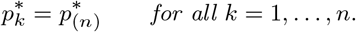

*That is, at an optimal division all subtasks have the same completion probability*.

*Proof* We prove the contrapositive. Suppose a given division is such that not all completion probabilities are equal. Then there exists a pair of consecutive subtasks, say *k* + 1 and *k*, with

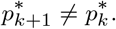

We will show that the division cannot be optimal.

Fix all subtasks other than *k* + 1 and *k*, and vary only the split of their combined cost

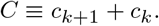

All terms in the total expected time *T* other than *T*_*k*+1_ + *T*_*k*_ remain unchanged, so it is enough to minimize

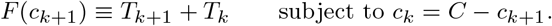

We first treat the case *k >* 1, so that both subtasks are interior subtasks.

As in Theorem 2, we define

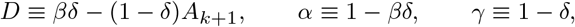

By Equation (22),

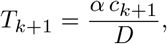

because the cumulative cost up to subtask *k* is

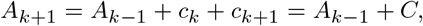

which is fixed. Likewise, for subtask *k* the cumulative cost is

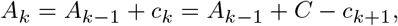

so

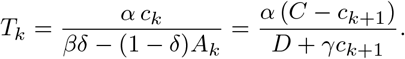

Therefore

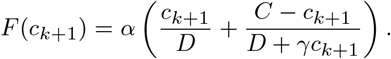

Differentiate with respect to *c*_*k*+1_:

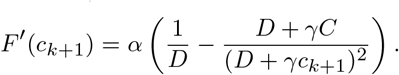

Hence the critical points satisfy

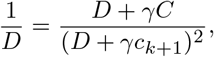

or equivalently

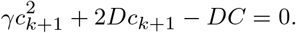

This quadratic has a unique solution in (0, *C*), since

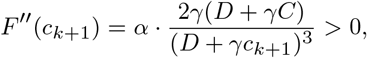

so *F* is strictly convex. Therefore this critical point is the unique minimizer.

Now compute the completion probabilities of these two subtasks. By Equation (34),

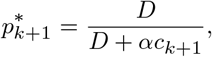

and for the next subtask,

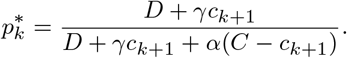

We compare them:

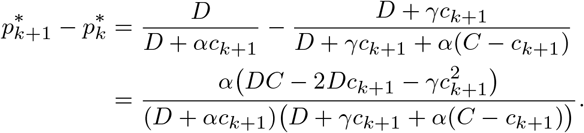

Therefore

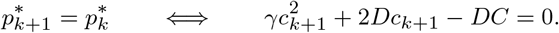

But this is exactly the first-order condition for minimizing *F*. Hence the unique minimizer of *T*_*k*+1_ + *T*_*k*_ is characterized by

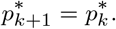

So if for some consecutive interior pair one has

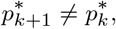

then that pair does not minimize its contribution to total expected time, and the whole division cannot be optimal.

It remains to handle the boundary pair (2, 1). Here subtask 1 is the last subtask, so by Equations (21) and (22),

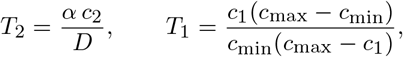

where now

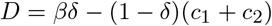

and *c*_1_ + *c*_2_ = *C* is fixed. Writing *c*_2_ = *C* − *c*_1_, we get

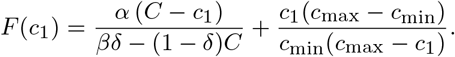

The derivative of *F* (*c*_1_) with respect to *c*_1_ is

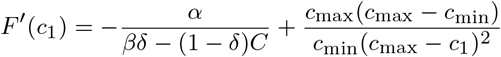

Using 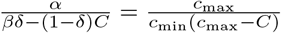, and Equations (18) and (19),

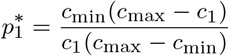

and

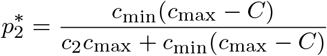

we can write

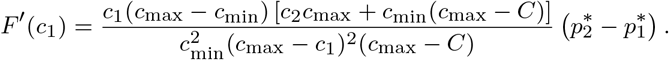

Hence,

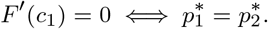

The function *F* is strictly convex since

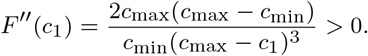

So, any interior critical point is the unique minimizer.

We have shown that for every consecutive pair of subtasks, the contribution of that pair to total expected time is minimized only when the two completion probabilities are equal. Therefore, if an entire division were optimal but had some consecutive pair with unequal probabilities, one could adjust only that pair and decrease total expected time, a contradiction.

Hence every consecutive pair in an optimal division must satisfy

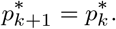

By transitivity,

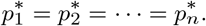

Denoting their common value by 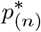 completes the proof.

### Theorem 4

(Optimal libertarian division assigns more cost to the second half) *Assume the libertarian regime and suppose subtask k is in the stochastic regime. Let subtask k, of cost c*_*k*_, *be divided into two consecutive subtasks b and a, with costs*

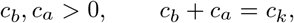

*where b precedes a*.

*If the split minimizes the expected time contributed by the two new subtasks, then*

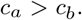

*Equivalently, the optimal split allocates more than half of the subtask cost to the second half*.

*Proof* We distinguish the cases *k* = 1 and *k >* 1.

**Case 1:** *k* = 1. Let *C* ≡ *c*_1_ + *c*_2_ denote the total cost of the two-subtask block after splitting the last subtask. By Equations (19) and (20),

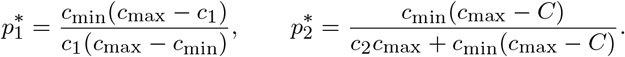

By Theorem 3, an optimal split equalizes the completion probabilities of the two halves. Therefore the minimizer satisfies 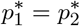.

Now compare the corresponding costs. Setting 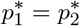 gives

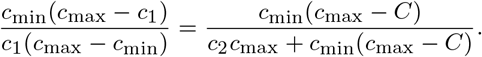

After cancellation and rearrangement,

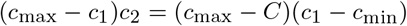

or equivalently,

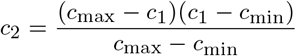

Hence

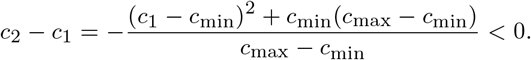

It follows that

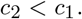

Thus, in the optimal split of the last subtask, the second half carries more cost than the first:

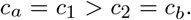

**Case 2:** *k >* 1.

As in the proof of Theorem 3, we consider a pair of consecutive subtasks, say *k* + 1 and *k* and define

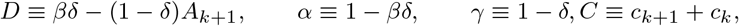

and consider

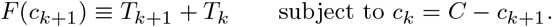

Using Equation (22),

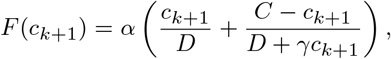

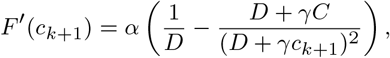

and the function *F* is strictly convex since

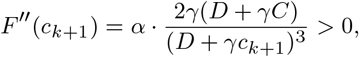

Evaluate the derivative at the midpoint in which 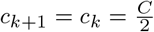:

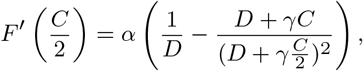

Multiplying by the positive quantity 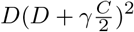,

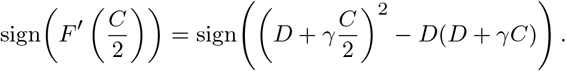

But

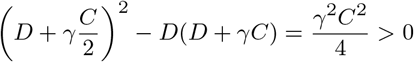

Hence

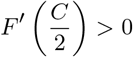

Because *F* is strictly convex and *F*′ *>* 0 is increasing at 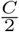, the minimum of *F* lies to the left of 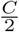. Therefore the minimizing split satisfies

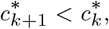

Together, in both cases, the optimal split assigns strictly more cost to the second subtask.

### Theorem 5

(Completion-time in the continuous libertarian case is Poisson distributed) *Assume the libertarian regime. Let the last subtask have cost c*_min_, *and divide the remaining cost c* − *c*_min_ *into subtasks of constant cost dc, where dc* → 0.

*Let T denote the completion time, and let N* = *T* − 1 *denote the number of pauses before completion. Then, in the limit dc* → 0,

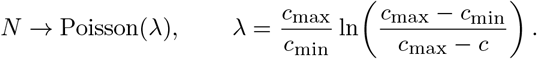

*Equivalently*,

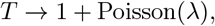

*In particular, the average time to task completion is* 1 + *λ and the probability that task is completed on the first day is e*^−*λ*^.

*Proof* For an interior subtask *k*, the probability that it will not be executed is given by Equation (20):

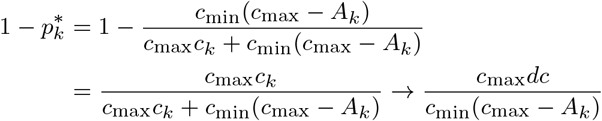

where we took the limit of *dc* → 0. So, over an interval of task-space length *dx*, the probability of a pause for cumulative cost *x > c*_min_ is *ν*(*x*)*dx* where

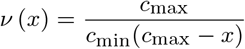

For *x < c*_min_ the task will be completed without any pauses. Thus, the pauses for *x > c*_min_ form an inhomogeneous Poisson process in task-space. The mean number of pauses up to total cost *c* is therefore,

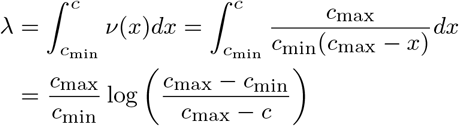

as claimed.

### Proposition 6

(Closed-form optimal stochastic sequence in the libertarian case) *Assume the libertarian regime and suppose the optimal division is stochastic, so that*

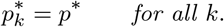

*Then, the optimal sequence of subtask costs has the form*

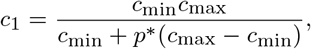

*and, for every k* ≥ 2,

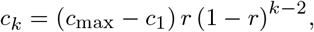

*where*

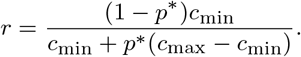

*Equivalently*,

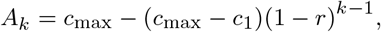

*where* 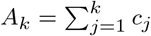.

*Proof* Since the optimal division is stochastic, all subtasks have the same completion probability *p*^∗^.

For the last subtask, Equation (19) gives

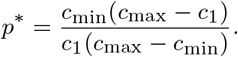

Solving for *c*_1_,

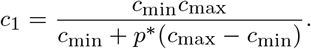

For every *k >* 1, Equation (20) gives

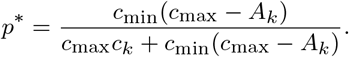

Rearranging,

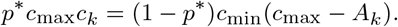

Since *A*_*k*_ = *A*_*k*−1_ + *c*_*k*_,

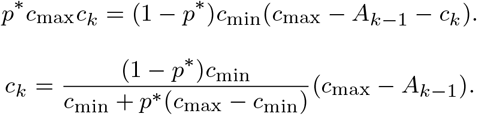

Define

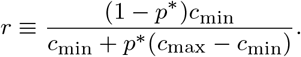

Then

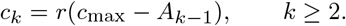

Therefore

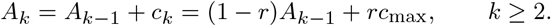

This linear recursion has solution

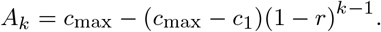

Finally,

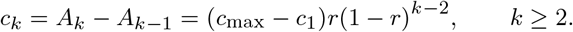

This proves the claimed closed form.

### Theorem 7

(Paternalist task splitting weakly reduces expected completion time) *Assume the paternalist regime and suppose subtask k is in the stochastic regime. Let subtask k, of cost c*_*k*_, *be divided into two consecutive subtasks b and a, with costs*

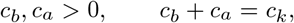

*where b precedes a. If both new subtasks remain in the stochastic regime, then*

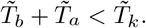

*Hence splitting a stochastic paternalist subtask strictly reduces the expected time contributed by that subtask, provided both resulting segments are stochastic*.

*Proof* By Equation (32),

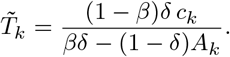

Define

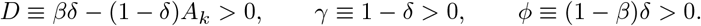

Then

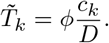

After splitting *c*_*k*_ = *c*_*b*_ + *c*_*a*_, the first new subtask *b* is evaluated at cumulative cost *A*_*k*_, whereas the second new subtask *a* is evaluated at cumulative cost *A*_*k*_ − *c*_*b*_. Therefore

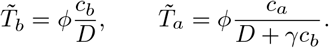

Since *c*_*b*_ *>* 0 and *γ >* 0,

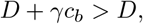

and hence

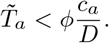

Adding,

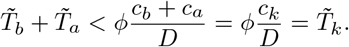

### Theorem 8

(Optimal paternalist division equalizes completion probabilities) *Assume the paternalist regime and suppose all subtasks are in the stochastic regime. Let a task of total cost c be divided into n sequential subtasks with costs c*_*n*_, …, *c*_1_, *and let* 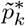 *denote the self-consistent probability of completing subtask k. If this division minimizes the expected time to task completion among all divisions with the same total cost c, then there exists a number* 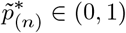 *such that*

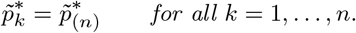

*That is, at an optimal paternalist division all subtasks have the same completion probability*.

*Proof* Suppose, toward a contradiction, that an optimal paternalist division has two consecutive subtasks with unequal completion probabilities. Let these be subtasks *k* and *k* − 1, and let

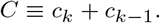

Because Equation (32) shows that 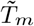 depends only on *A*_*m*_ and *c*_*m*_, redistributing the fixed block cost *C* between the pair (*k, k* − 1) leaves all other subtask times unchanged. Thus, to minimize the total expected completion time, the division must minimize the contribution of this pair.

Define, as before,

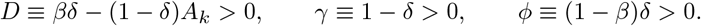

Writing *c*_*k*−1_ = *C* − *c*_*k*_, Equation (32) gives

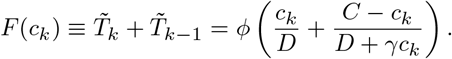

Differentiating *F* (*c*_*k*_) with respect to *c*_*k*_,

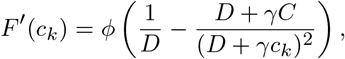

and

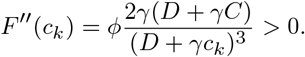

Hence *F* is strictly convex, so its unique minimizer is characterized by *F*′(*c*_*k*_) = 0, namely

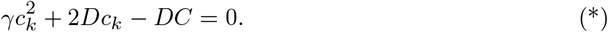

On the other hand, using Equation (31),

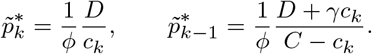

Therefore

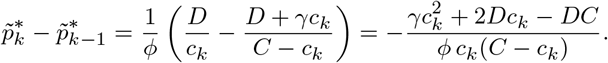

Thus condition (∗) is equivalent to

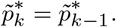

So the unique repartition of the block cost *C* that minimizes the contribution of the pair (*k, k*− 1) is exactly the repartition for which the two completion probabilities are equal. If an allegedly optimal paternalist division had a pair of consecutive subtasks with unequal probabilities, repartitioning their combined cost would strictly reduce total expected completion time, contradicting optimality.

Hence every consecutive pair in an optimal paternalist division must satisfy

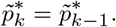

By transitivity,

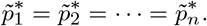

### Proposition 9

(Closed-form optimal stochastic sequence in the paternalist case) *Assume the paternalist regime and suppose the optimal division is stochastic, so that*

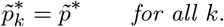

*Then the optimal sequence of subtask costs is geometric:*

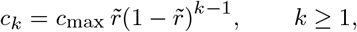

*where*

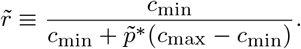

*Equivalently*,

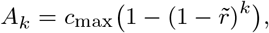

*where* 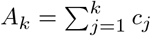.

*Proof* Since the optimal paternalist division is stochastic, all subtasks have the same completion probability 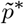.

Equation (31) gives

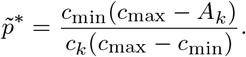

Using *A*_*k*_ = *A*_*k*−1_ + *c*_*k*_,

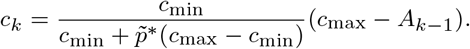

Define

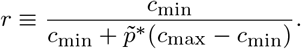

Then

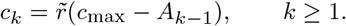

It follows that

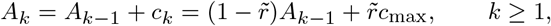

with *A*_0_ = 0. This linear recursion has solution

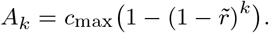

Finally,

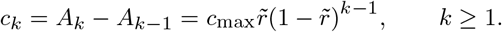

This proves the claimed closed form.

### Theorem 10

(Optimal paternalist division assigns more cost to the second half) *Let a paternalist subtask of cost C be divided into two consecutive subtasks b and a, with*

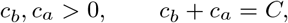

*where b precedes a. Then exactly one of the following holds*.

1. *If there exists a split for which both resulting subtasks are deterministic, then every such split is optimal and*

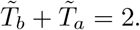
2. *If no split makes both resulting subtasks deterministic, then the unique time-minimizing split is stochastic for both subtasks and satisfies*

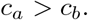

*Proof* Define

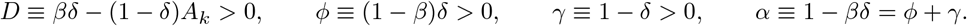

For a paternalist split, the expected completion time is

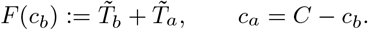

Using Equation (32), in the stochastic regime,

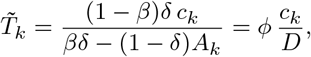

Thus, the first subtask *b* is deterministic iff

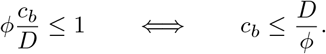

Similarly, the second subtask *a* is deterministic iff

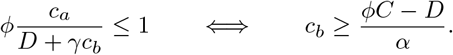

Therefore both subtasks are deterministic iff

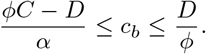

If such a split exists, then

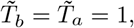

so

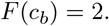

Since at most one action is allowed per day, any two-subtask split must take at least two days, so 2 is the global minimum. Hence every split satisfying these conditions on *c*_*b*_ is optimal.

Now assume that no split makes both subtasks deterministic. Then

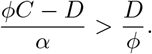

Set

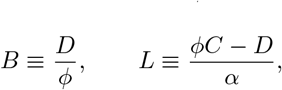

so that *B < L*.

On the interval 0 *< c*_*b*_ ≤ *B*, subtask *b* is deterministic and subtask *a* is stochastic, so

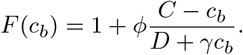

Differentiating,

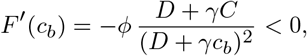

so *F* is strictly decreasing on (0, *B*].

On the interval *L* ≤ *c*_*b*_ *< C*, subtask *a* is deterministic and subtask *b* is stochastic, so

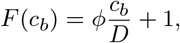

and therefore

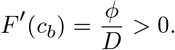

Thus *F* is strictly increasing on [*L, C*).

Finally, on the interval *B < c*_*b*_ *< L*, both subtasks are stochastic, and

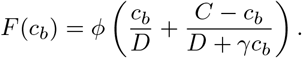

Differentiating,

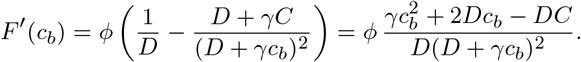

Hence critical points satisfy

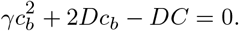

whose positive solution is

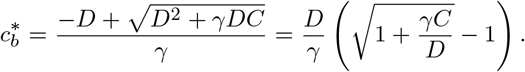

Because

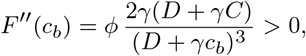

this solution is the global minimum.

Using 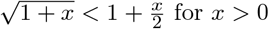 for *x >* 0, we obtain

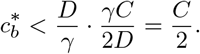

Therefore

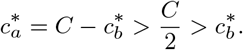

So, when no deterministic–deterministic split is available, the unique optimal split assigns more work to the second half.

### Theorem 11

(Paternalist division reduces procrastination relative to the same libertarian division) *Consider a task divided into N >* 1 *sequential subtasks with cumulative costs*

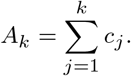

*Assume the stochastic self-consistent policy in both the libertarian and paternalist models. Let T*_*k*_ *and* 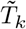 *denote the expected time required to complete subtask k in the libertarian and paternalist regimes*.

*Then for k >* 1,

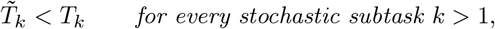

*and therefore the expected total completion time satisfies*

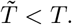

*Proof* From Equation (22), in the libertarian case,

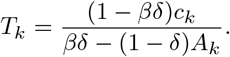

From Equation (32) in the paternalist case,

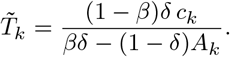

Thus

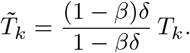

Since 0 *< β <* 1 and 0 *< δ <* 1, we have

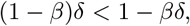

and hence

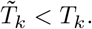

Summing over subtasks yields 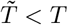.

## Footnotes

1 As is customary now, ‘hyperbolic’ will refer to any discount function that exhibits decreasing impatience (Prelec 1989), e.g., present bias in the quasi-hyperbolic model (Laibson 1997).

2 O’Donoghue and Rabin (2001) note the existence of such a strategy in footnote 14 but exclude it from their analysis without computing it. Their description of the strategy as “Pareto-efficient” uses a restricted notion of efficiency, internal to the set of equilibrium strategies; we return to this distinction below.

3 The term “sophisticated” is standard in the procrastination literature (Strotz 1955; O’Donoghue and Rabin 2001), where it often suggests deliberate forecasting of future selves’ behavior. Our usage is weaker. The agent’s beliefs are accurate at equilibrium, but she need not reach that equilibrium through explicit forecasting. As we show in Section 2.4, the stochastic policy can arise through myopic reinforcement on local feedback. “Sophisticated” should therefore be read as an unbiased-equilibrium condition, not as a cognitive achievement.

4 Throughout the manuscript, we number the tasks backwards, so that the last task is denoted by 1, the penultimate task by 2, and so on.

5 If no interior fixed point exists, the dynamics converge to the deterministic boundary policy.

6 This is the sense in which our usage differs from O’Donoghue and Rabin (2001)’s footnote 14. If only equilibrium strategies are allowed as comparators, the mixed strategy is trivially efficient because no self has a strict deviation incentive at the equilibrium probability. We instead use the unrestricted Pareto criterion across temporal selves, following the multi-self welfare framework of Bernheim and Rangel (2009).

## Notes

### Competing Interest Statement

The authors have declared no competing interest.

